# Synthetic repertoires derived from convalescent COVID-19 patients enable discovery of SARS-CoV-2 neutralizing antibodies and a novel quaternary binding modality

**DOI:** 10.1101/2021.04.07.438849

**Authors:** Jule Goike, Ching-Lin Hsieh, Andrew Horton, Elizabeth C. Gardner, Foteini Bartzoka, Nianshuang Wang, Kamyab Javanmardi, Andrew Herbert, Shawn Abbassi, Rebecca Renberg, Michael J. Johanson, Jose A. Cardona, Thomas Segall-Shapiro, Ling Zhou, Ruth H. Nissly, Abhinay Gontu, Michelle Byrom, Andre C. Maranhao, Anna M. Battenhouse, Varun Gejji, Laura Soto-Sierra, Emma R. Foster, Susan L. Woodard, Zivko L. Nikolov, Jason Lavinder, Will N. Voss, Ankur Annapareddy, Gregory C. Ippolito, Andrew D. Ellington, Edward M. Marcotte, Ilya J. Finkelstein, Randall A. Hughes, James M. Musser, Suresh V. Kuchipudi, Vivek Kapur, George Georgiou, John M. Dye, Daniel R. Boutz, Jason S. McLellan, Jimmy D. Gollihar

## Abstract

The ongoing evolution of SARS-CoV-2 into more easily transmissible and infectious variants has sparked concern over the continued effectiveness of existing therapeutic antibodies and vaccines. Hence, together with increased genomic surveillance, methods to rapidly develop and assess effective interventions are critically needed. Here we report the discovery of SARS-CoV-2 neutralizing antibodies isolated from COVID-19 patients using a high-throughput platform. Antibodies were identified from unpaired donor B-cell and serum repertoires using yeast surface display, proteomics, and public light chain screening. Cryo-EM and functional characterization of the antibodies identified N3-1, an antibody that binds avidly (K_d,app_ = 68 pM) to the receptor binding domain (RBD) of the spike protein and robustly neutralizes the virus *in vitro*. This antibody likely binds all three RBDs of the trimeric spike protein with a single IgG. Importantly, N3-1 equivalently binds spike proteins from emerging SARS-CoV-2 variants of concern, neutralizes UK variant B.1.1.7, and binds SARS-CoV spike with nanomolar affinity. Taken together, the strategies described herein will prove broadly applicable in interrogating adaptive immunity and developing rapid response biological countermeasures to emerging pathogens.

## Introduction

The rapid global dissemination of the severe acute respiratory syndrome coronavirus 2 (SARS-CoV-2)^1^, the cause of coronavirus disease 19 (COVID-19)^2^, has highlighted our extreme vulnerability to novel microbial threats. The speed of SARS-CoV-2 transmission and absence of widespread adaptive immunity created a pandemic that overwhelmed the international medical community. This situation was exacerbated by the scarcity of treatment options, especially early in the pandemic. Functional immune repertoire analysis has the potential to efficiently address this scarcity. By analyzing primary immune responses directed towards emerging pathogens, newly elicited antibodies can be identified and rapidly deployed to treat patients.

The COVID-19 pandemic has stimulated global research efforts to identify SARS-CoV-2-neutralizing antibodies for therapeutic and prophylactic applications. Initial attempts to identify neutralizing antibodies focused on screening extant antibodies elicited against previous SARS-CoV strains. These efforts were largely unsuccessful owing to limited cross-reactivity^3^, primarily due to SARS-CoV-2’s sequence divergence in the receptor-binding domain (RBD) of its trimeric spike protein^4^.

The ectodomain (ECD) of the SARS-CoV-2 spike (S) protein is essential for initial binding and subsequent entry of the virus into human cells and has been the primary target for therapeutics and vaccine formulations. The ECD consists of the S1 subunit, containing an N-terminal domain (NTD) and RBD, and the S2 subunit, containing the fusion machinery that mediates entry into host cells. The RBD initiates attachment through interaction with angiotensin-converting enzyme 2 (ACE2)^5–7^. The functional significance of the ACE2-RBD interaction makes the RBD a prime target for neutralizing antibodies^8–10^. Targeting this domain increases the selective pressure on the RBD, which may promote the emergence of escape mutants that maintain virulence. Indeed, neutralizing antibodies targeting a single epitope can induce virus escape in cell culture, quickly rendering antibodies ineffective^11^.

The FDA approved antibody cocktail of REGN10933 and REGN10987^12^ targets distinct epitopes of the RBD and sustains some neutralization activity against new SARS-CoV-2 variants B.1.1.7 and B.1.351, originally identified in the UK and South Africa, respectively^13–15^. Recently, a deep mutational scan of the RBD found that the single amino acid mutation E406W increased the IC50 of REGN10933 by more than 300-fold and also detrimentally affected REGN10987 binding^16^. The scan also identified amino acid changes that compromised the epitope of Eli Lilly’s antibody LY-CoV016^16^. Moreover, the mass sequencing of 5,085 SARS-CoV-2 genomes from the Houston metropolitan area published in September 2020 already identified numerous spike protein mutations that affect the existing neutralizing antibodies’ abilities to bind to their epitopes^17^.

The B.1.1.7 (UK), B.1.1.248 (Brazil), and B.1.351 (South Africa) variants are of special concern because they have each accumulated multiple spike protein mutations. These mutations are located in the NTD, RBD, and furin cleavage site. NTD-directed antibodies in particular show reduced or abolished binding to these strains^14^. More concerning than the loss of binding ability, however, is the reduced neutralizing activity of convalescent plasma from patients infected in the spring of 2020^14, 18^. Furthermore, currently available vaccines are largely based on an earlier prefusion-stabilized spike variant (Wuhan). When assayed, sera from patients vaccinated with the Moderna and Pfizer/BioNTech vaccines showed a significant decrease in neutralization activity towards strain B.1.351, while B.1.1.7 was only mildly affected^14^. It is therefore critical to identify neutralizing antibodies to a broad spectrum of non-overlapping epitopes on the spike protein to achieve highly potent and persistent neutralization^11^.

To this end, we developed a multi-pronged antibody discovery and informatics strategy involving both proteomic analysis of donor sera and selection of combinatorially paired heavy and light chain (VH-VL) libraries from donor B-cell receptor repertoires. Specifically, we used Ig-Seq proteomics to identify candidate variable heavy chains (VHs) of serum antibodies binding SARS-CoV-2 RBD or ECD. We identified productive light chain pairs for these heavy chains either from a bioinformatically derived set of “public light chains” (PLCs) or through combinatorial pairing with donor light chain repertoires via yeast surface display (YSD) selection. In addition to Ig-Seq interrogation of circulating IgGs, we employed YSD for selection of high-affinity antibodies by combinatorial display of all donor VH-VL pairs as Fab fragments through multiple rounds of selection.

Together, this holistic strategy allowed us to probe the secreted circulating antibody repertoire and the nascent cellular repertoire of a primary immune response. By interrogating the combined repertoire, we gained valuable insight into the antibody response elicited by SARS-CoV-2 and discovered neutralizing antibodies to multiple distinct domains of the spike protein. We report the structural characterization of two such neutralizers, including a highly potent neutralizing antibody (N3-1) that binds a quaternary epitope of the trimeric spike protein via a novel binding modality. Finally, we tested binding to emerging spike variants of SARS-CoV-2 and demonstrated N3-1 maintains robust binding to these variants and neutralizes B.1.1.7.

## Results

### Ig-Seq analysis of the serological repertoire reveals candidate heavy chains

Ig-Seq provides a proteomic snapshot of the serum antibody repertoire by using mass spectrometry to identify heavy chain complementarity-determining region 3 (CDR3) peptides of antigen-enriched antibodies (**Fig. 1**)^19^. To identify relevant SARS-CoV-2 antibodies present in the serological repertoire, we first isolated antibodies from the serum of two donors by antigen enrichment chromatography using immobilized SARS-CoV-2 RBD and ECD fragments. We then employed Ig-Seq mass spectrometry to identify abundant IgG clonotypes. A total of 15 and 21 unique clonotypes were identified with high confidence from donors 1 and 2, respectively (**Extended Data Figs. 1-2**). This is notably lower than what has been observed in previous Ig-Seq studies^19–22^, perhaps because the previous studies involved donors after boost vaccination which is likely to elicit a more robust and diverse response than the primary response to natural SARS-CoV-2 infection observed here. Regardless, the clonotypes identified provided a sufficient set of VH candidates to express and characterize as recombinant anti-SARS-CoV-2 monoclonal antibodies (mAbs) in combination with suitable light chains.

**Figure 1.**
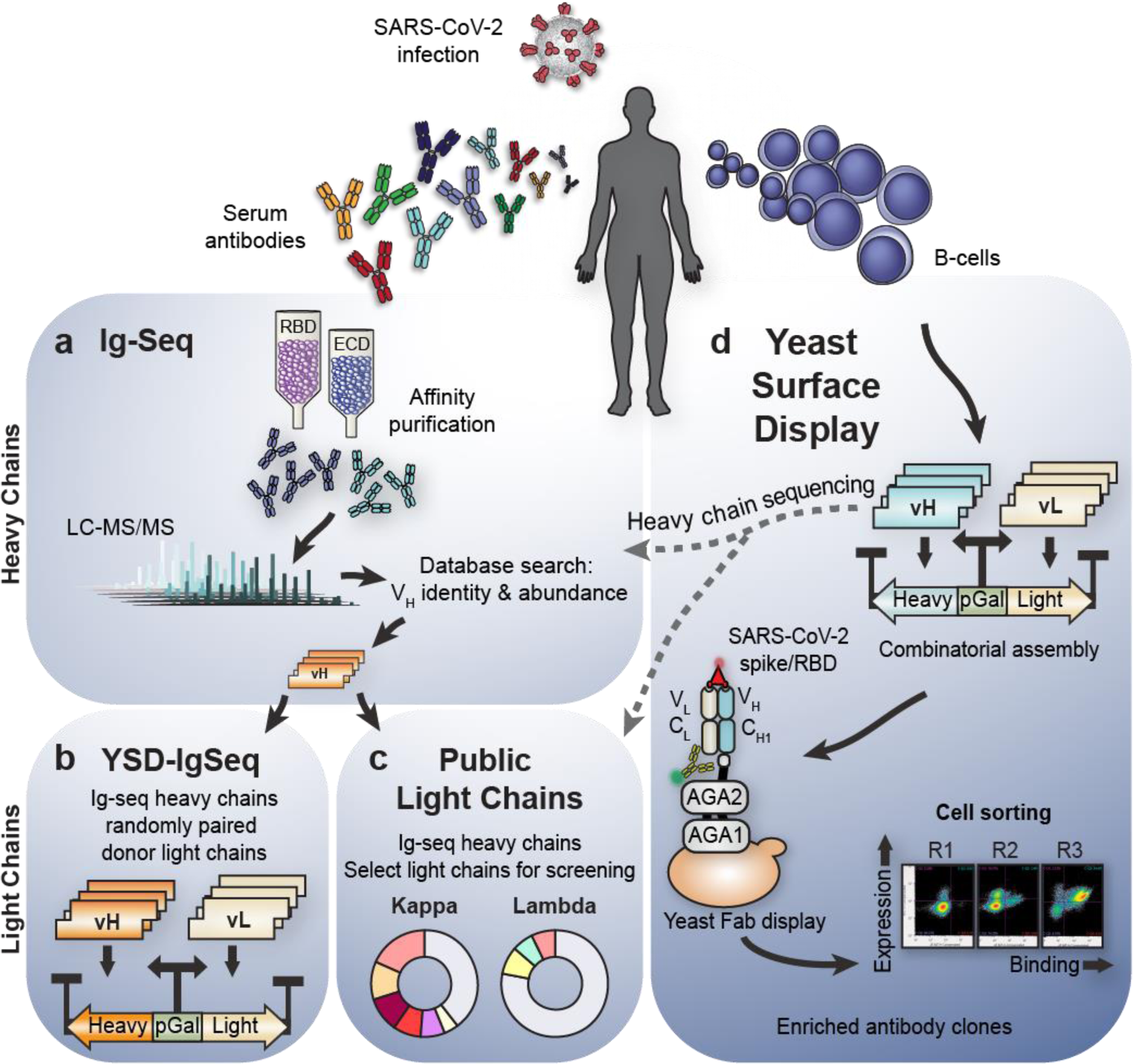
Overview of complementary strategies for discovering convalescent patient-derived anti-SARS-CoV-2 neutralizing antibodies. Serum antibodies and B-cells are isolated from patient blood post infection. **a**, IgG is antigen-enriched, digested, and analyzed with tandem mass spectrometry. Searching against a database of donor B cell receptor sequences yields antigen-specific heavy chain candidates. **b**, To find productive VH-VL pairs, the proteomically identified VH are randomly combined with donor VL, generating a large library of Fab amplicons for selection in yeast. **c**, Additional productive VH-VL are recovered by matching each proteomic VH candidate against a fixed set of nine public light chains significantly overrepresented in prior repertoire studies. All combinations are screened for binding. **d**, Yeast surface display provides an independent method to recover VH or VHVL candidates. VH and VL sequences are amplified from donor B-cells, and pairs are randomly assembled into large Fab libraries and selected using fluorescent cell sorting. Yeast pools are then sequenced after each selection to track individual clone dynamics round over round. Promising candidates are expressed as IgG for further characterization.

### Public light chains recover productive pairs for most VHs

While Ig-Seq provides valuable information for antibody discovery, it does not identify light chain partners, which usually requires additional laborious techniques. To expedite light chain discovery, we analysed a published dataset of natively paired memory B-cell sequences^19^ from three healthy donors and bioinformatically derived a panel of nine light chains that show high frequencies of productive pairings with a diverse set of VH sequences (**Fig. 1c**).

At the time of blood draw, the donors in the dataset were healthy and asymptomatic. This lack of polarization and relative homeostasis potentially affords broader insight into common VH-VL pairings, independent of antigen specificity. Indeed, many of the observed light chains paired promiscuously with a diverse set of VH genes. The V gene usage of undiscerning light chains was remarkably consistent among the three donors and therefore were termed “public light chains” (PLCs). From our secondary analysis of this dataset, we identified six kappa V genes that accounted for 61% of the kappa VH-VL pairs and three lambda V genes representing 22% of lambda pairs (**Fig. 2a**). Together, these nine public light chain (PLC) V genes accounted for 43% of observed VH-VL pairs in the analyzed repertoires.

**Figure 2.**
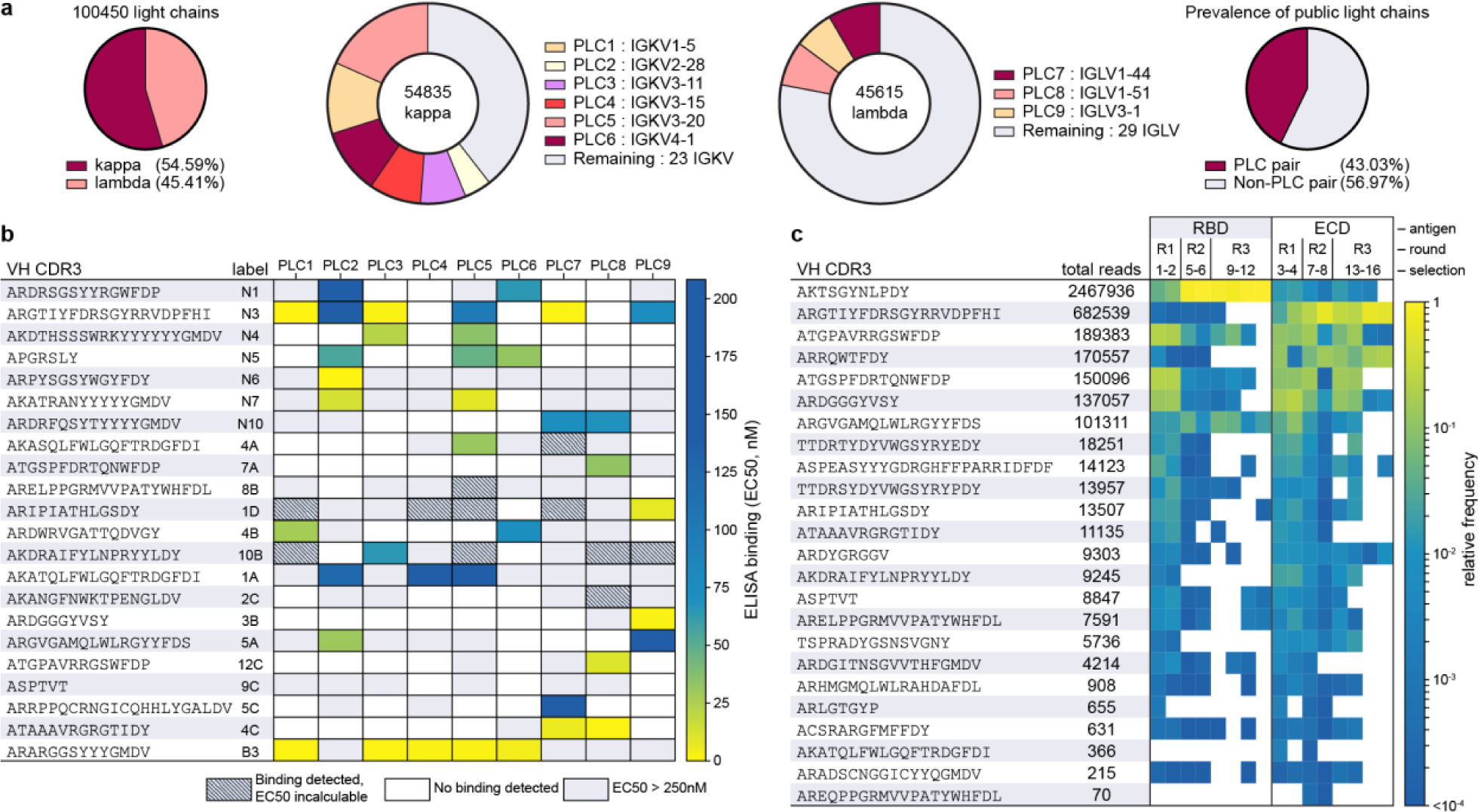
Public light chain and YSD-IgSeq abundance and screening. **a**, Nine PLCs emerge from analysis of 100,450 previously published paired VH-VL sequences. **b**, Screening with VHs (H-CDR3 depicted) identified by Ig-Seq and YSD against the panel of nine PLCs to determine productive VHVL pairings. IgG mAbs ELISA EC50s revealed that partnering VHs with PLCs can create low nM affinity binders. Grey boxes indicate that binding was detected but at an affinity too low to determine an EC50 in the concentration range tested. White boxes, no binding was detected. **c**, VH CDR3 read counts and relative abundance from the YSD-IgSeq experiment shown for MinION reads of 1.7 kb to 2.1 kb. Heatmap values are CDR3 frequencies, or read counts normalized within each respective sample.

We hypothesized that this abundance could indicate an enhanced ability to form productive VH-VL pairs independent of VH sequence and antigen specificity. To test this hypothesis, we constructed germline versions of these light chains with their most frequently observed J gene as published on the iRepertoire website^20–23^ (**Extended Data Table 1**) and obtained full-length light chains for each PLC. SARS-CoV-2 specific Ig-Seq heavy chains were then expressed with each PLC as a full-length IgG1. EC50 determination via ELISA using recombinant spike protein demonstrated that of the 22 VHs tested, all paired productively with one or more of the PLCs, resulting in several highly potent and specific antibodies (**Fig. 2b**). Through this simple and rapid screening method we identified a panel of functional antibody candidates for SARS-CoV-2 neutralization testing.

### Yeast Surface Display-IgSeq identifies additional candidate antibodies

Our PLC screening strategy provides a method to rapidly generate functionally paired antibodies, but the decreased sequence space of the nine PLCs may limit the ability to efficiently identify potent binders. Thus, to search for additional high-affinity antibodies, we randomly paired 24 Ig-Seq heavy chains with donor-derived light chain libraries for yeast surface display (YSD) selections (**Fig. 1b**), which allowed us to efficiently screen Fabs for each heavy chain/light chain library combination. In YSD, heavy/light chain pairs were displayed as Fabs on the surface of a humanized yeast strain that expresses human protein chaperones to improve antibody expression. We then selected antigen-binding Fabs using two-color fluorescence cell sorting to maximize Fab expression and binding to either SARS-CoV-2 spike (S-2P) or RBD (**Extended Data Figs. 3-4**). As Fab enrichment over successive rounds of selection indicates antigen specificity, we performed three rounds and identified and quantified productive VH-VL pairs based on MinION nanopore sequencing of Fab amplicons with sequence error correction and complementary Illumina reads (**Fig. 2c**, **Extended Data Figs. 5-6**). When light chain preference was compared, we observed extensive correspondence between YSD selection and PLC screening.

### Parallel selections *via* yeast surface display yield functionally diverse antibodies

Much of the early stage primary antibody response is inaccessible to Ig-Seq discovery, as detection is limited to secreted antibody proteins with sufficient abundance in serum^24^. To broaden our search for neutralizing antibodies, we employed our YSD platform to directly mine the cellular repertoire for high affinity B-cell receptor sequences (**Fig. 1d**). We expressed and displayed large VH-VL libraries (>10^7^) created by combinatorial pairing of donor IgG heavy and light chain amplicons. To maximize the functional diversity of selected antibodies, we performed three rounds of selection on the initial libraries, using various gating strategies to ensure phenotypic diversity (i.e. a variety of antibody expression and antigen binding strengths). Ultimately, the parallelized selection strategy yielded populations with distinct binding characteristics as confirmed by cytometric phenotyping of single clones (**Extended Data Figs. 4, 7**). We observed progressive enrichment of specific antibodies with the dominant clone from each course of selections generally approaching 20% of the total population after round three (**Extended Data Fig. 8**). Individual clones from enriched populations were selected as candidates for further characterization either at random or based on their abundance from the bioinformatic analysis. Importantly, YSD detected enrichment of VH sequences that were not observed by Ig-Seq proteomics, demonstrating the advantages and increased sensitivity of this multi-pronged approach to antibody discovery.

### Non-cognate VH-VL pairs are potent neutralizing antibodies to SARS-CoV-2

We initially screened the isolated IgG against the ECD, NTD, and RBD of the spike protein using ELISA to identify the most promising candidates for authentic live virus neutralization of SARS-CoV-2. We prioritized non-RBD antibodies in our screens in order to gain insights into alternative neutralization mechanisms. From binding data of the full-length IgGs, we selected 94 candidate VH-VL pairs for live virus neutralization, and 37 pairs successfully neutralized authentic SARS-CoV-2/WA1 (**Fig. 3a, Extended Data Table 2**). Curve fitting calculated IC50 values as low as 18 pM, with seven antibodies showing sub-nanomolar values (8-32, 18 pM; N3-1, 251.9 pM; 8-131, 300 pM; 8-114, 350 pM; 12C8, 836 pM; A7V3, 955 pM; and 7-6, 977 pM). Twenty-three neutralizing antibodies had IC50 values less than 100 nM (**Fig. 3**). Our high affinity NTD VH-VL pairs (8-32, 8-131, 8-114, 12C8, A7V3, 7-6, 4C7, 4C8, 7A8) showed strong preference for IGHV1-24 gene usage. PLC screening successfully identified VL partners for both YSD and Ig-Seq derived VHs (**Fig. 3b-e**). Seven of the mAbs were shown to be directed to the S2 domain. N3-1 possesses a highly potent RBD-directed IGHV4-31 heavy chain and displayed strong neutralization activity with several PLCs as well as YSD VLs (**Extended Data Table 2**).

**Figure 3.**
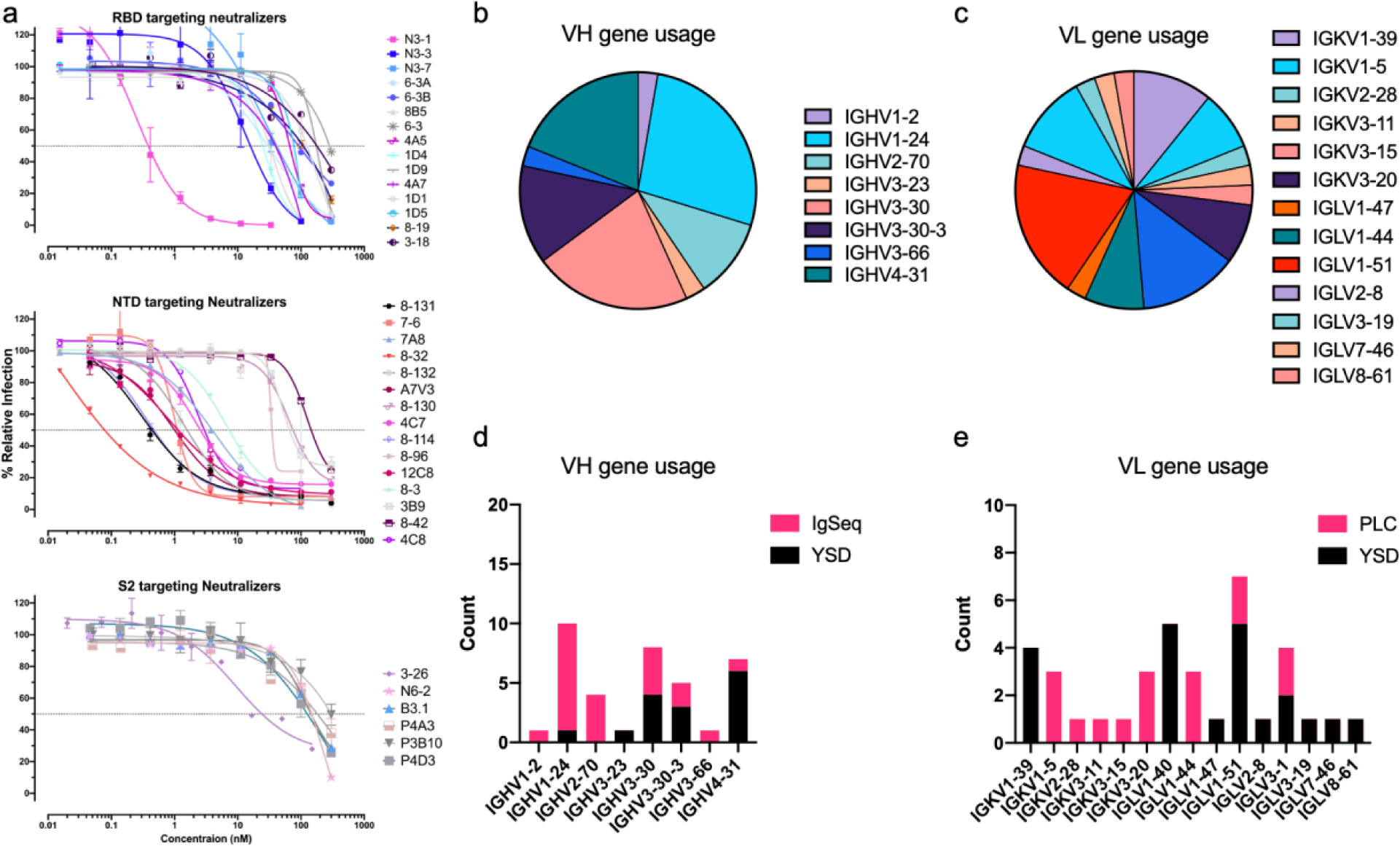
Non-cognate paired antibodies neutralize SARS-CoV-2 live virus. **a**, SARS-CoV-2 WA1 live virus neutralization assays. The graphs depict the neutralizers binned by their target spike subdomain. **b**, VH gene usage of the neutralizing antibodies as a fraction of the whole. **c**, Stacked bar chart showing neutralizer VH gene usage broken down by discovery method. **d-e**, Same analysis described in b-c but by VL gene.

### Common binding mode of an NTD-directed antibody

A7V3 was of interest because of the different donor origins of the VH and VL. Its IGHV1-24 was discovered in both IgSeq and YSD from donor 1. However, the IGLV1-51 was mapped to donor 2 and is also related to PLC8. The VH of A7V3 showed a strong preference to PLC8 in ELISA screening, and related light chains appear in six other neutralizing VH-VL pairs, commonly pairing with IGHV1-24 VHs. To ascertain its potential binding mode and contributions from both variable regions, we determined a cryo-EM structure of A7V3 Fab complexed with SARS-CoV-2 S-ECD. The initial 3D reconstruction from 715,398 particles had a global resolution of 3.0 Å (**Fig. 4a**). Although the map revealed an A7V3 Fab molecule bound to each NTD of the trimeric spike, only one Fab defined clear density. Further 3D classification was performed to resolve potential heterogeneity in the particle set. One class was obtained consisting of one fourth of the total particles and exhibited defined density for an NTD and its bound Fab. A local refinement of this map focused on the NTD and Fab yielded a reconstruction with a well-resolved binding interface (**Fig. 4b**). Similar to the first structurally defined NTD-targeted antibody 4A8^25^, the heavy chain of A7V3 plays a dominant role in binding the N3 loop (residues 141-156) and N5 loop (residues 246-260) of the NTD. The light chain of A7V3 makes limited contact with the NTD, and is therefore likely to play a structural role in positioning the heavy chain. H-CDR1, H-CDR3, and L-CDR2 collectively form a concave surface packed tightly against the N5 loop. Specifically, Trp100d in H-CDR3, and both Tyr49 and Pro55 in L-CDR2, form a hydrophobic cage to enclose Pro251 at the tip of the N5 loop (**Fig. 4c**). Glu31 from H-CDR1 is expected to form a salt bridge with Arg246, and Asp101 from H-CDR3 is expected to form a hydrogen bond with the hydroxyl group of Tyr248. Pro97 and Phe98 from H-CDR3 insert into a groove flanked by the N3 and N5 loops (**Fig. 4d**). As opposed to the major binding interface on the N5 loop, A7V3 only makes a minor contact with the N3 loop, mainly mediated by H-CDR1 and H-CDR2. Akin to a conserved Phe from H-CDR2 of 4A8, Phe51 contacts Lys147 via a π-cation interaction (**Fig. 4d-e**). In stark contrast, H-CDR3 of A7V3 is much shorter than its counterpart in 4A8, which contains a Phe that forms a π-π stacking interaction with Trp152 on the N3 loop. Collectively, the structural characterization of A7V3 demonstrates that our semi-synthetic approach is capable of yielding a novel neutralizing antibody that binds to the NTD using an established modality.

**Figure 4.**
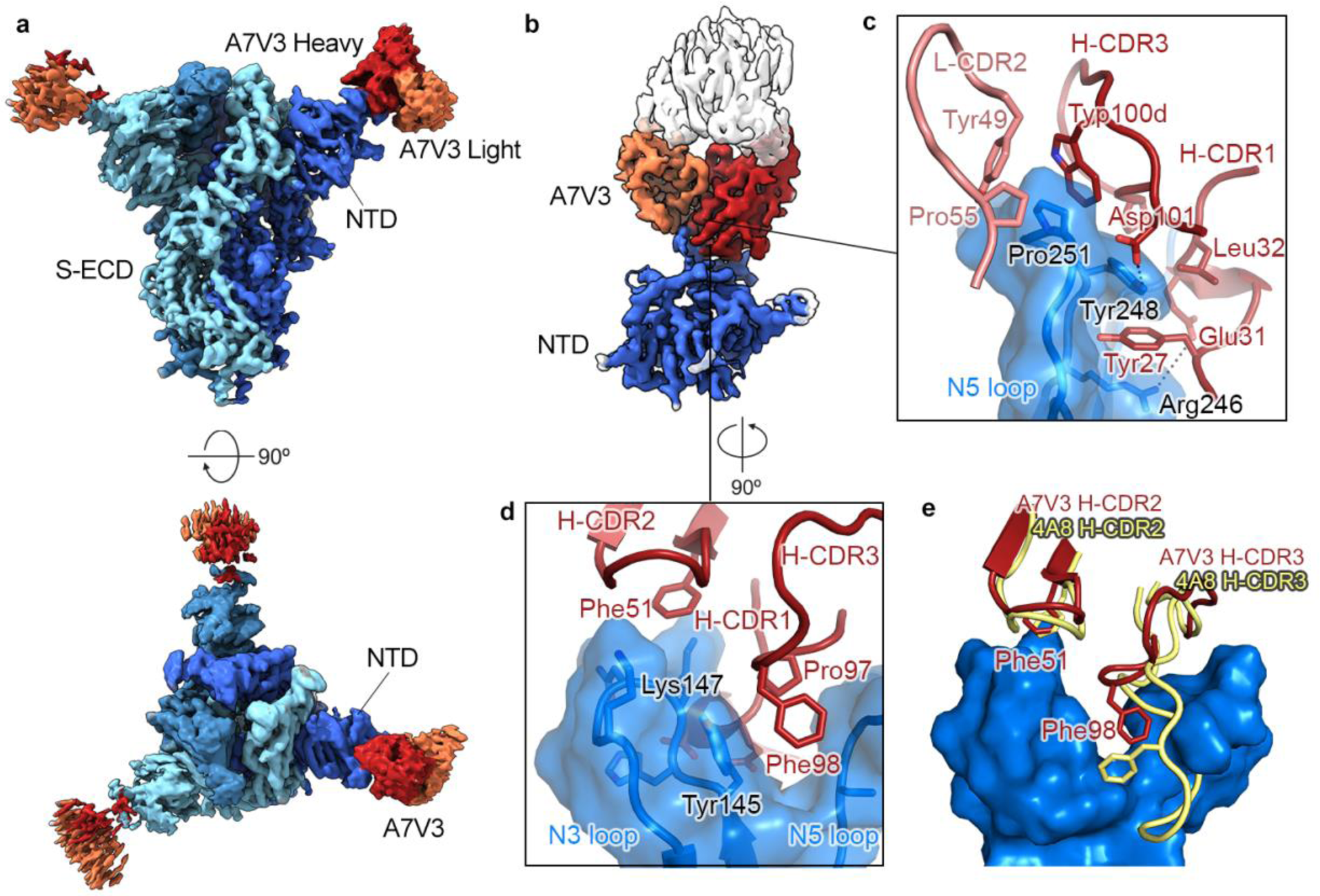
NTD-directed mAb A7V3 exhibits a common binding mode. **a**, Cryo-EM structure of A7V3 bound to SARS-CoV-2 S at a global resolution of 3.0 Å. Side view and top-down views of the complex are shown in the upper and the lower panel, respectively. Each protomer is depicted in steel blue, royal blue and sky blue. The heavy chain of A7V3 is colored firebrick, and the light chain is colored coral. **b**, Focused map of A7V3 bound to NTD reveals a major binding site on the N3 and N5 loops. **c**, The tip of the N5 loop is surrounded by the hydrophobic residues from L-CDR2 and H-CDR3. Arg246 and Tyr248 contact Glu31 and Asp101 via polar interactions. **d**, Pro97 and Phe98 in H-CDR3 insert into a groove walled by the N3 and N5 loops. Conserved Phe51 in H-CDR2 forms a pi-cation interaction with Lys147 **e**, Superimposed structure of 4A8-NTD complex (PDB ID: 7C2L) with A7V3-bound NTD. The molecular surface of the N3 and N5 loops is shown in blue. Unlike 4A8, the relatively short H-CDR3 from A7V3 barely contacts the N3 loop.

### An RBD-targeted antibody exhibits a novel quaternary binding mode

Structurally defined RBD-targeted antibodies or nanobodies complexed with SARS-CoV-2 S may be generally grouped into two types based on binding sites. The first type, mainly discovered from the IGHV3-53 germline family, recognizes the ACE2-binding site and directly blocks host receptor engagement (e.g. antibodies CC12.1^26^, C105^27^ and VHH E^28^). The second type, from a more diverse germline family, recognizes a relatively conserved region on the RBD that is mostly buried and contacts a neighboring RBD in the down conformation (e.g. C135^29^, S309^30^ and VHH V^28^). From negative stain EM analysis of the IgG N3-1-spike complex, we observed density for two Fabs, likely derived from the same IgG molecule, bound to multiple RBDs on a single trimeric spike (**Extended Data Figs. 12**). We also found the affinity of N3-1 IgG to SARS-CoV-2 spike is nearly 1000-fold stronger than N3-1 Fab to spike (**Extended Data Fig. 13**).

To investigate this unique binding mode, we determined a cryo-EM structure of N3-1 Fab bound to SARS-CoV-2 S-ECD to a global resolution of 2.8 Å (**Fig. 5a**). We observed that two N3-1 Fabs bind to a single trimeric spike with one Fab binding to RBD in the up conformation and the other Fab simultaneously engaging two RBDs: one in the up conformation and one in the down conformation. We performed focused refinement on the Fab bound to the two RBDs (**Fig. 5b**), which substantially improved the interpretability of the map in this region. The well-resolved Fab-RBD binding interface revealed two completely distinct epitopes on RBD-up and RBD-down (**Fig. 5c**). For RBD-up interaction, contacts are made by the Fab H-CDR1, H-CDR3 and L-CDR2, which together bury 862 Å^2^ surface area. For the RBD-down interaction, contacts are made by the Fab H-CDR2, H-CDR3 and L-CDR3, which buries 696 Å^2^ surface area. Notably, the relatively long H-CDR3 loop (18 a.a.) engages both RBDs via hydrophobic and polar interactions. H-CDR3 residues Tyr98, Phe99 and Arg100a pack against a hydrophobic pocket formed by Tyr369, Phe377, Lys378 and Pro384 on RBD-up, which are highly conserved between SARS-CoV and SARS-CoV-2. This pocket is barely exposed when RBD is in the down conformation and is part of the shared epitopes targeted by cross-reactive antibodies CR3022 and COVA1-16^31^. Lys378, from the upper ridge of the pocket, forms a cation-π interaction with Tyr98 of H-CDR3, and its amine group is expected to form a salt bridge with Asp100h of H-CDR3. In addition, main chain atoms of Cys379 and Tyr369 form hydrogen bonds with Phe99 and Arg100a from H-CDR3, strengthening this primary Fab binding interface on RBD-up. Furthermore, the sidechain guanidinium of Arg408 on RBD-up likely has polar interactions with Tyr49 and Glu55 from L-CDR2, which along with H-CDR1, constitutes the secondary binding interface on RBD-up.

**Figure 5.**
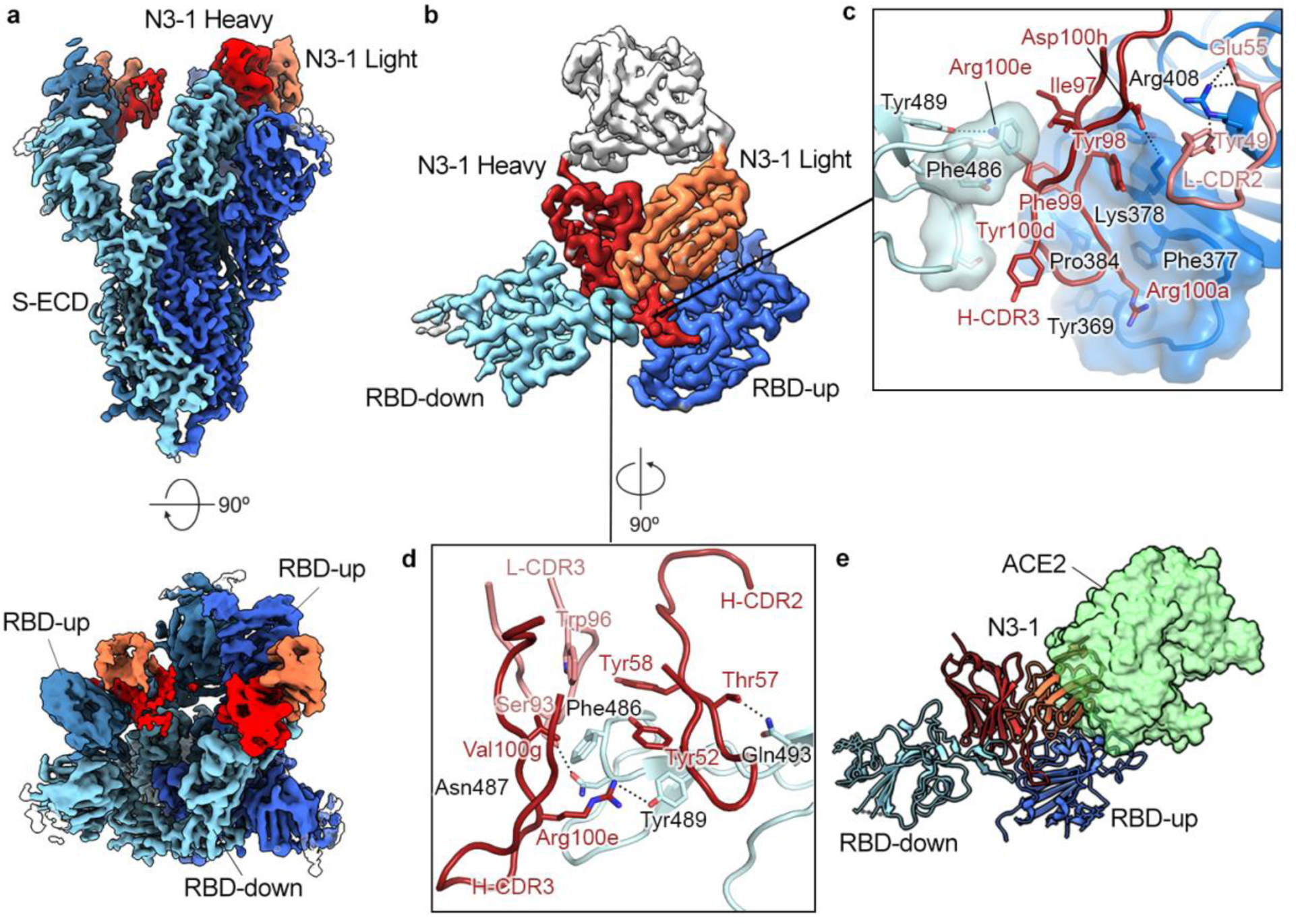
RBD-directed mAb N3-1 exhibits a unique binding mode by recognizing two distinct epitopes. **a**, Cryo-EM structure of N3-1-bound SARS-CoV-2 spike at a global resolution of 2.8 Å. Side view and top-down views of the complex are shown in the upper and the lower panel, respectively. Each protomer is depicted in steel blue, royal blue and sky blue. The heavy chain of N3-1 is colored firebrick, and the light chain is colored coral. **b**, Focused map of N3-1 bound to RBDs in the up and down conformations with five CDRs involved in the binding interface. **c**, One face of H-CDR3 contacts a conserved hydrophobic pocket (transparent royal blue surface) on RBD-up. Arg408 forms multiple polar interactions with Tyr49 and Glu55. The other face of H-CDR3 contacts the ACE2-binding site on RBD-down. H-CDR1 contacts the epitopes on RBD-up, but it is omitted for clarity. **d**, The epitope on RBD-down is centered on Phe486, which fits into a hydrophobic surface formed by Trp96, Tyr58, Tyr52, Arg100e and Val100g (clockwise). Arg100e also forms a cation-pi interaction with Phe486 and a hydrogen bond with Tyr489. **e**, Superimposed crystal structure of RBD-ACE2 complex (PDB ID: 6M0J) with N3-1 bound RBDs. The molecular surface of ACE2 is shown in transparent pale green. The light chain of N3-1 heavily clashes with ACE2. The ACE2-binding site on RBD-down is completely blocked by H-CDR2, H-CDR3 and L-CDR3 [**d**].

In contrast, the N3-1 binding site on RBD-down overlaps with the ACE2-binding site, where 11 of 19 N3-1 epitope residues are also involved in ACE2 binding (**Fig. 5d**). Phe486 on RBD-down inserts into a hydrophobic pocket formed by Trp96 (L-CDR3), Tyr52, Tyr58 (H-CDR2), Arg100e and Val100g (H-CDR3). The sidechain guanidinium of Arg100e not only forms a hydrogen bond with Tyr489, but also contacts Phe486 through cation-π interactions. In addition, Asn487 and Gln 493 on RBD-down are expected to form polar interactions with Ser93 of L-CDR3 and Thr57 of H-CDR2. The angle of approach of N3-1 prevents ACE2 binding by both trapping one RBD in the down position, thereby preventing exposure of the ACE2 binding site, as well as by sterically inhibiting ACE2 access to the bound ‘up’ RBD. (**Fig. 5e**). Collectively, the N3-1 antibody engages an extensive quaternary epitope on neighboring RBDs through a novel binding modality.

### mAb N3-1 binds robustly to spike proteins from emerging variants of concern

Next, we tested whether binding of A7V3 (NTD-targeting) or N3-1 (RBD-targeting) was detrimentally affected by mutations emerging in SARS-CoV-2 variants of concern (VOC). We assessed binding to the 12 most abundant SARS-CoV-2 variants from Houston^17^ and did not observe detrimental losses in binding avidity (**Extended Data Table 5**). Next, we used mammalian surface display of spike variants to test binding of N3-1 to mutations that were previously shown to reduce or abrogate binding to one or both of the Regeneron antibodies (REGN10987 and REG10933) as well as a commonly found mutation in Australia (S477N). Strikingly, N3-1 was able to bind each of the mutant spike variants. N3-1 binding was slightly enhanced by 10 of the 12 mutant spike proteins tested (L455A, G476D, E406W, Q493F, K444A, K444Q, V445A, G446A, G446V, and S477N) using this assay. Variant Y453A was indistinguishable from binding to the prefusion stabilized SARS-CoV-2 S D614G, and F486K was the only mutation shown to marginally reduce binding (∼6%) (**Extended Data Fig. 15**). We then tested binding to B.1.1.7, B.1.1.248, and B.1.351 spike protein variants using the same mammalian surface display assay. Binding of A7V3 was reduced by all three variants, a result predicted from our structural work. In contrast, N3-1 binding was largely unperturbed by the accumulated mutations (**Fig. 6a**) of B.1.1.7, B.1.1.248, and B.1.351 spike proteins in this assay (**Fig. 6b**). To validate the mammalian display screening results, we examined the binding of the SARS-CoV-2 S Wuhan-Hu-1, and its variants B.1.1.7, and B.1.351 to N3-1 Fab or N3-1 IgG using surface plasmon resonance (SPR). N3-1 Fab binding to each of the variants was only able to be fit to a heterogenous binding model (**Extended Data Fig. 13**), consistent with the structural studies. N3-1 IgG exhibited comparable binding to Wuhan-Hu-1 and its variants, with 67.9 pM apparent affinity to Wuhan-Hu-1, 296 pM apparent affinity to B.1.1.7, and 300 pM apparent affinity to B.1.351 (**Fig. 6c-e**). We also tested binding via SPR to SARS-CoV spike protein and discovered that N3-1 IgG cross-reacts with SARS-CoV spike with 28 nM affinity (**Extended Data Figure 13**). Finally, we corroborated these findings with live virus neutralization assays with circulating VOC B.1.1.7. We found that the wildtype WA/1 strain was fully neutralized by both N3-1 and A7V3, but B.1.1.7 was only neutralized by N3-1 at the same titer as WA/1 (**Fig. 6f**), consistent with our structure-based predictions.

**Figure 6.**
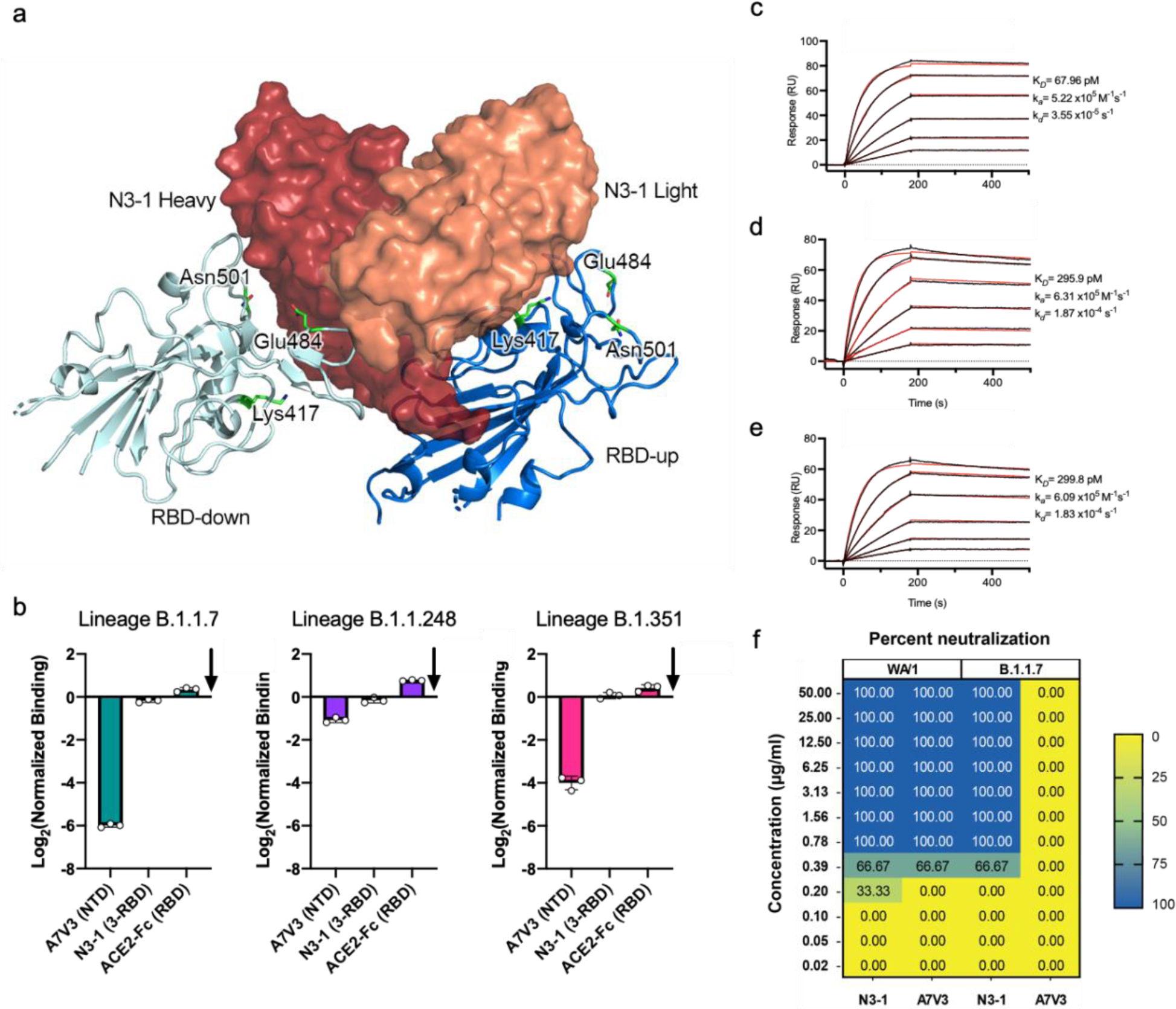
N3-1 binds to circulating spike variants. **a**, Model of N3-1 bound to RBD-up and RBD-down. Mutation sites in B.1.1.7, B.1.1.248, and B.1.351 are highlighted in green. **b**, The B.1.1.7, B1.1.248, and B.1.351 spike variants were tested with A7V3, N3-1, and chimeric human ACE2-Fc using mammalian display of spike variants and compared to SARS-CoV-2 Wuhan-Hu-1 S-HexaPro-D614G (baseline arrow). **c-e**, Binding of IgG N3-1 to SARS-CoV-2-S WuHan-Hu-1 **[c]**, variants B.1.1.7 **[d]** and B.1.351 **[e]** were assessed by surface plasmon resonance using an NTA sensor chip. Binding data are shown as black lines. For **[c-e]**, the best fit was achieved using a 1:1 binding model and shown as red lines. **f**, Neutralization of B.1.1.7 and WA/1 by N3-1 and A7V3, where neutralization is measured as the percent of replicates showing complete neutralization at a given concentration (see methods).

## Discussion

SARS-CoV-2 and its evolved progeny represent a moving target for therapeutic development. While the rapid response of the global research community has led to the discovery of many neutralizing mAbs^25, 32, 33^ the emergence and widespread dissemination of viral variants such as the U.K. variant B.1.1.7, Brazilian variant B.1.1.248, and South African variant B.1.351 stresses the need for continued efforts to rapidly identify and develop antibodies that can bind to diverse epitopes. By combining proteomic analyses of antibody repertoires from patient samples and yeast-display with public light chain pairing, we developed a high-throughput antibody discovery strategy that identified 37 neutralizing antibodies that target multiple regions of the SARS-CoV-2 spike protein, including the RBD, NTD, and the S2 domain, and that broadly bind multiple viral variants.

To accelerate the discovery of neutralizing antibodies, we first performed Ig-Seq proteomic analyses on patient samples. However, because Ig-Seq does not provide information on relative affinities, epitope targeting, and light chain pairings, we adopted two strategies for light chain discovery: a rapid screening method utilizing public light chains (PLCs) and a YSD light chain selection. Ultimately, we found that both methods independently yielded multiple neutralizing antibodies, with several of the most potently neutralizing mAbs converging in two or more of our approaches.

The neutralizing antibodies identified relied on eight IGHV genes, including IGHV3-30 and IGHV1-2, consistent with studies that show these genes are overrepresented in SARS-CoV-2 neutralizing antibodies^32^. In addition, IGHV1-24 was previously found to contribute to a number of NTD-directed neutralizing antibodies^25, 34^ and was observed in 10 of the neutralizing antibodies herein. We found representative neutralizers from each of our discovery strategies, although each method yielded one or more unique IGVH or IGVL genes, validating the high coverage afforded by this multi-pronged approach.

The rapid PLC screening method appears particularly useful not only for expediting functional testing of Ig-Seq-derived heavy chains, but also to generate high-affinity antibodies. For instance, N3-1 exhibits a sub-nanomolar IC-50 (252 pM) and binding affinity (68 pM), while A7V3 has an IC-50 of 950 pM. A7V3 is a particularly interesting demonstration of the utility of the strategy, because its heavy chain was first identified from one donor, while the light chain was mapped to another donor. Initial PLC screening indicated a nearly identical IGLC VJ gene (PLC8) as a productive partner, which further confirmed the utility of PLC screening for productive antibody VH-VL pairings.

We resolved high-resolution cryo-EM structures of these two newly discovered neutralizing antibodies, and found that N3-1 targets the RBD, whereas A7V3 targets the NTD. Structural analysis of N3-1 revealed that the L-CDR2 and L-CDR3s were both engaged in binding, which suggests that our bioinformatically derived PLCs may directly contribute to neutralization. A7V3 bound to a common neutralization site on the NTD^34^, and ultimately, PLC8 was found to productively pair with three different neutralizing antibodies that were most commonly associated with NTD-binding heavy chains. Our work not only establishes the first empirical demonstration that public light chains can form functional pairings across heavy chains, but also debuts their utility as a prolific method for high affinity antibody discovery, potentially improving rapid-response antibody discovery for neo-antigens in future epidemics. Our results further suggest that naturally paired repertoires may be unnecessary for efficient antibody discovery, in contrast to a recently published work where cognate pairing was deemed necessary for effective neutralization^35^.

The cryo-EM structure of N3-1 is of particular importance because it reveals a novel binding mechanism at a quaternary epitope of the spike trimer, thus allowing this antibody to bind the RBD in both the ‘up’ and ‘down’ conformations. This greatly enhances the binding affinity of the full mAb as it enables binding to all three RBD subunits in the spike protein trimer. As a consequence of this unique mechanism of action, N3-1 is minimally perturbed by spike mutations in the newly emerged variants B.1.1.7, B.1.1.248, and B.1.351, where it was shown to still have low picomolar affinities and neutralized B.1.1.7 with a similar titer to WA/1. N3-1 is also capable of binding the distantly related SARS-CoV spike protein with low nanomolar affinity (28 nM). Given the mechanism of binding and the fact that the N3-1 binding epitope for the RBD-down conformation corresponds to the ACE2-binding site, we fully expect N3-1 to show neutralization of VOCs not tested here.

The strategy we described here will aid in the rapid identification and development of monoclonal antibodies to extant SARS-CoV-2 variants, and can be readily utilized for identifying antibody therapeutics for variants yet to be discovered and for other explosive pathogen zoonoses. Candidate mAbs may be produced in less than 10 days, and have desirable characteristics – presence in immune responses, public light chain robustness, validation of expression via yeast display – that may enable production at scale. Because of the modular nature of the synthetic biology-like discovery process, the overall strategy could also be readily extended to affinity maturation or directed evolution of tighter binding mAbs to distinct antigens or epitopes. In the long term, the ability to store, combine, and continually reassess repertoires should prove broadly useful in not only discovering novel antibodies with potential therapeutic applications, but also for studying the evolution of adaptive immunity.

## Online Methods

### Strains and media

Yeast strain EBY100 (MATa AGA1::GAL1-AGA1::URA3 ura3-52 trp1 leu2-delta200 his3-delta200 pep4::HIS3 prbd1.6R can1 GAL) was acquired from ATCC (cat. no. MYA-4941) and used for antibody expression and selection. To improve antibody expression the human chaperones BIP (binding immunoglobulin protein) and PDI (protein disulfide isomerase) were genomically integrated as an expression cassette in the HO locus. Yeast were grown in rich medium (YPD; Takara, cat. no. 630409) or in selective medium for leucine prototrophs after library transformation (Takara cat. no. 630310). YEP-galactose was used for expression of displayed antibody libraries (1% yeast extract, 1% bacto-peptone, 0.5% NaCl, 2% galactose, 0.2% glucose).

### Antigens and antibodies

Spike antigen was biotinylated using the EZ-link kit (Thermo Scientific, cat. no. 21435) and labeled with streptavidin-AF647 (Invitrogen, cat. no. S32357). RBD was labeled with a mouse anti-human Fc-AF647 (Southern Biotech, cat. no. 9042-31). Fab library light chains were labeled with anti-FLAG M2-FITC (Sigma, cat. no. F4049).

### Donors

Blood was collected from 3 PCR-confirmed, symptomatic patients. Donors 1 and 2 were collected at day 12 post-onset of symptoms. Donor 3 was collected on day 11. None of the donors were hospitalized or experienced severe disease. PBMCs and plasma were both collected by density gradient centrifugation using Histopaque-1077 (Sigma-Aldrich).

### Preparation of serum antibodies for Ig-Seq proteomic analysis

Serum samples were prepared for Ig-Seq analysis as previously described^34^. Briefly, total IgG was isolated from plasma using Pierce Protein G Plus Agarose (Pierce Thermo Fisher Scientific) and cleaved into F(ab’)2 fragments with IdeS protease. Antigen-specific F(ab’)2 was enriched by affinity chromatography against recombinant SARS-CoV-2 S-2P or RBD protein cross-linked to NHS-activated agarose resin (Thermo Fisher Scientific). Eluted F(ab’)2 fractions were concentrated by vacuum centrifugation and prepared for mass spectrometry-based proteomic analysis as previously described.^24^

### LC-MS/MS analysis of antigen-enriched antibodies

Liquid chromatography-tandem mass spectrometry analysis was carried out on a Dionex Ultimate 3000 RSLCnano system coupled to an Orbitrap Fusion Lumos Mass Spectrometer (Thermo Scientific). Samples were loaded onto an Acclaim PepMap 100 trap column (75 μm × 2 cm; Thermo Scientific) and separated on an Acclaim PepMap RSLC C18 column (75 μm × 25 cm; Thermo Scientific) with a 3%-40% acetonitrile gradient over 60 min at a flow-rate of 300 nl/min. Peptides were eluted directly into the Lumos mass spectrometer using a nano-electrospray source. Mass spectra were acquired in data-dependent mode with a 3 sec. cycle time. Full (MS1) scans were collected by FTMS at 120,000 resolution (375-1600 m/z, AGC target = 5E5). Parent ions with positive charge state of 2-6 and minimum intensity of 3.4E4 were isolated by quadrupole (1 m/z isolation window) and fragmented by HCD (stepped collision energy = 30+/-3%). Fragmentation (MS2) scans collected by ITMS (rapid scan rate, AGC target = 1E4). Selected ions and related isotopes were dynamically excluded for 20 sec (mass tolerance = +/-10ppm).

### Antibody variable chain sequencing

Peripheral blood mononuclear cells (PBMCs) from processed donor samples were provided in Trizol as a kind gift from Dr. Gregory C. Ippolito^34^. IgG and IgM VH and VL cDNA libraries were separately amplified from PBMC RNA of three donors and sequenced to create donor specific reference databases, from which the complete amino acid sequences of serum IgG proteins could subsequently be determined based on their mass spectral identifications. **Extended data Figure 1** shows the distribution and diversity of clonotypes and V-gene usage for the variable heavy chain repertoires of donors 1 and 2.

### Ig-Seq MS data analysis

Mass spectra were analyzed using Proteome Discoverer 2.2 software (Thermo Scientific). Precursor masses were first recalibrated with the Spectrum File RC node using a consensus human reference proteome database (UniProt) with common contaminants (MaxQuant) and precursor mass tolerance of 20 ppm. Recalibrated mass spectra were searched against a custom database for each donor consisting of donor-derived VH sequences, VL sequences, and the human and contaminant sequences using the Sequest HT node. Mass tolerances of 5 ppm (precursor) and 0.6 Da (fragment) were used. Static carbamidomethylation of cysteine (+57.021 Da) and dynamic oxidation of methionine (+15.995 Da) were considered. False discovery rates for peptide-spectrum matches (PSMs) were estimated by decoy-based error modelling through the Percolator node. Label-free quantitation (LFQ) abundances were calculated from precursor areas using the Minora Feature Detector and Precursor Ions Quantifier nodes.

Resulting PSMs were filtered according to methods described (Boutz, 2014). Briefly, peptide sequences differing only by isoleucine/leucine substitution were considered equivalent and combined into a single PSM. PSMs were re-ranked by posterior error probability, q-value, and Xscore. Only top-ranked, high-confidence PSMs (FDR < 1%) were retained for each scan. If two or more PSMs had identical top-ranked scores, they were considered ambiguous and removed. PSMs for the same peptide sequence were summed and the average mass deviation (AMD) was calculated for each peptide. Peptides with AMD greater than 2 ppm were filtered out. Peptides mapping to VH sequences from a single clono-group were considered clono-specific. Clono-specific peptides overlapping the CDR3 sequence by four amino acids or more were considered CDR3-informative.

For each clono-group, PSMs and LFQ abundances of clono-specific CDR3-informative peptides were summed. Ratios of elution:flow-through PSMs and LFQ abundances were calculated; only clono-groups with both ratios > 5 were considered elution-specific.

### Library assembly and bacterial transformation

Donor B-cell VH and VL amplicons were amplified via PCR to include adapters for cloning into yeast expression vectors. Assembly into the yeast kappa and lambda expression vectors was done via Golden Gate assembly. Library assemblies were prepared in 20 μL reactions as follows: 2 μL 10X AARI buffer (ThermoFisher Scientific, cat. no. B27), 0.4 μL 50X oligo buffer (ThermoFisher Scientific, cat. no. ER1582), 0.2 μL 100 mM ATP (ThermoFisher Scientific, cat. no. R0441), 20 fmol backbone DNA, 40 fmol VH and VL amplicons, 0.5 μL (2 U/μL) AARI endonuclease (ThermoFisher Scientific, cat. no. ER1582), and 0.5 μL T7 ligase (3000 U/μL) (NEB, cat. no. M0318). Each assembly was scaled up to 16 total reactions in 8-well strips. Thermocycling consisted of the following protocol: 37°C, 15 minutes; 37°C, 2 minutes, 16°C, 1 minute; go to step 2, x74; 37°C, 60 minutes; 80°C, 15 minutes; hold at 4°C. Assemblies were consolidated and column purified using Promega Binding Solution (Promega, cat. no. A9303) to bind DNA to a Zymo-spin II column (Zymo Research, cat. no. C1008). The column was washed twice with DNA Wash Buffer (Zymo Research, cat. no. D4003) and eluted in 30 μL nuclease-free water. For library transformations, DH10B cells were diluted 1:100 from confluent culture into 50 mL Superior broth (AthenaES, cat. no. 0105). When cells reached an OD_600_ of 0.4-0.6, they were washed 3X with cold 10% glycerol and resuspended to a final volume of 600 μL. The purified library was added to cells and electroporated at 1.8 kV in an E. coli Pulser electroporator (Bio-Rad) using Genepulser 0.2 cm cuvettes (Bio-Rad, cat. no. 1652086) at 200 μL per transformation.

### Library transformation into yeast and protein expression

Purified libraries were linearized for integration into the yeast genome via homologous recombination at the Leu2 locus. For each 1 μg library plasmid, 0.5 μL NotI (10 units/μL) (NEB, cat. no. R0189) was used with the supplied Buffer 3.1 in 10 μL. Reactions were incubated at 37°C overnight and heat inactivated at 80°C for 20 minutes. Digests were pooled and column purified as described in previous sections and eluted in 25 μL nuclease-free water. Our strain was electroporated as described elsewhere^1^. We found that 10 μg of linearized DNA was sufficient for library sizes of 10^6^, and that library sizes could reach >10^7^ with 20 μg DNA. Transformed yeast were recovered in SD -Leu medium (see “strains and media” section). Libraries were passaged once at 1:100 before protein expression to reduce contamination from untransformed cells. To express Fab libraries, yeast were washed in YEP-galactose (see “strains and media”) and diluted 1:10 into 10 mL final volume. Cells were induced for 48 hours at 20°C with shaking.

### Fab library labeling and selection

Expressed yeast libraries were harvested at 100 μL (representing approximately 10^7^ cells) and washed with PBSA buffer (1X PBS, 2 mM EDTA, 0.1% Tween-20, 1% BSA, pH 7.4). Antigen was incubated with cells in 1 mL in PBSA at 200 nM at RT for one hour, washed with PBSA at 4C, and labeled with secondary antibodies (mouse anti-human FITC, 1:100; streptavidin-AF647, 1:100; mouse anti-human Fc-AF647, 1:50). Cells were washed 2X and resuspended in 2 mL in cold PBSA for sorting. Cell sorting was performed using a Sony SH800 fluorescent cell sorter. For first round libraries, 10^7^ events were sorted into 2 mL SD-Leu medium supplemented with penicillin/streptomycin (Gibco, cat. no. 15140122). Cells were recovered by shaking incubation for 1-2 days for further rounds of selection or plated directly for phenotyping clones.

### Next generation sequencing

Genome extraction was performed on yeast cultures of libraries and sorted rounds underwent genome extraction using a commercial kit (Promega, cat. no. A1120) with zymolyase (Zymo Research, cat. no. E1004). 100 ng genomic template was used to amplify the heavy and light chains separately or as one amplicon for short or long-read sequencing, respectively. For amplification of heavy chain genes only, primers JG.VHVLK.F and JG.VH.R were used. For amplification of light chain genes only, primers JG.VL.F and JG.VHVK.R or JG.VHVL.R were used for kappa and lambda vectors, respectively. For amplification of paired genes, primers JG.VHVLK.F and JG.VHVK.R or JG.VHVL.R were used. Amplicons were column purified and deep sequenced with an iSeq. In parallel, we obtained ∼1.8 kb sequences spanning the entire VH and VL using MinION nanopore sequencing (Oxford Nanopore Technologies Ltd., MinION R10.3).

### Colony PCR and Sanger sequencing

Sorted yeast populations were plated on SD -Leu and 8-32 colonies per plate were picked into 2 mL microplates either by hand or using a QPIX 420 (Molecular Devices) automatic colony picker. Cultures were grown at 1000 rpm at 3 mm orbit at 30°C overnight. Cells (20 μL) were transferred to a fresh microplate and washed with 1 mL TE buffer (10 mM Tris, 1 mM EDTA). Cells were incubated with 20 μL zymolyase solution (5 mg/mL zymolyase, 100T in TE) at 37°C for 1 hour. Cells (5 μL) were then used in colony PCR to amplify the paired heavy and light chains. Amplicons were column purified with the Wizard SV 96 PCR Clean-Up System (Promega, cat. no. A9342) and yields were quantified with a Nanodrop spectrophotometer or the Quant-it Broad-Range dsDNA kit (Invitrogen, cat. no. Q33130). Approximately 10 ng (2.5-5 μL) of purified PCR products were then subjected to Sanger sequencing.

### Long-read sequencing (donors 1 & 2)

Sequencing libraries were prepared from 18 amplicon samples using the Native Barcoding Kit (Oxford Nanopore Technologies; cat. no. EXP-NBD103) paired with the Ligation Sequencing Kit (Oxford Nanopore Technologies; cat. no. SQK-LSK109) according to the manufacturer’s directions. Between four and eight sequencing libraries per flow cell were pooled for sequencing on three MinION flow cells (Oxford Nanopore Technologies; R9.4.1) for 72 hours on an Oxford Nanopore Technologies MinION Mk1B device (Oxford Nanopore Technologies). Raw data was basecalled using the high accuracy model in Guppy (v.3.5.2).

### Short-read sequencing (phenotyping plates)

Sequencing libraries were prepared from 308 amplicon samples using the Nextera DNA Flex Library Preparation kit (Illumina; cat. no. 20018705) according to the manufacturer’s directions. Sequencing libraries were pooled and sequenced (2×151bp) on an iSeq 100 (Illumina; California, USA) using iSeq 100 i1 Reagents v.1 (Illumina; cat. no. 20021533).

### Long-read sequencing (donor 3)

Sequencing libraries were prepared from 32 amplicon samples using the Native Barcoding Kit (Oxford Nanopore Technologies; cat. no. EXP-NBD104) paired with the Ligation Sequencing Kit (Oxford Nanopore Technologies; cat. no. SQK-LSK109) according to the manufacturer’s directions. Between five and eight sequencing libraries per flow cell were pooled for sequencing on five GridION flow cells (Oxford Nanopore Technologies; R9.4.1) for 72 hours on a GridION Mk1 device (Oxford Nanopore Technologies; Oxford, England, UK). Raw data was live basecalled using the high accuracy model in Guppy (v.3.2.10).

### Short-read sequencing (donor 3)

Sequencing libraries were prepared from 32 samples using the Nextera DNA Flex Library Preparation kit (Illumina; cat. no. 20018705) according to the manufacturer’s directions. Sequencing libraries were pooled and sequenced (2×151bp) on an iSeq 100 (Illumina; California, USA) using the iSeq 100 i1 Reagents v2 (Illumina; cat. no. 2009584).

### Long-read sequencing (YSD-IgSeq)

Sequencing libraries were prepared from 16 amplicon samples using the Native Barcoding Kit (Oxford Nanopore Technologies; Cat. No. EXP-NBD104) paired with the Ligation Sequencing Kit (Oxford Nanopore Technologies; Cat. No. SQK-LSK109) according to the manufacturer’s directions. Four sequencing libraries were pooled per flow cell and sequenced on four GridION flow cells (Oxford Nanopore Technologies; R9.4.1) for 72 hours on a GridION Mk1 device (Oxford Nanopore Technologies; Oxford, England, UK). Raw data was live basecalled using the high accuracy model in Guppy (v.4.0.11).

### Short-read sequencing (YSD-IgSeq)

Sequencing libraries were prepared from 16 samples using the Nextera DNA Flex Library Preparation kit (Illumina; cat. no. 20018705) according to the manufacturer’s directions. Sequencing libraries were pooled and sequenced (2×150 bp) on an iSeq 100 (Illumina; California, USA) using the iSeq 100 i1 Reagents v2 (Illumina; Cat. No. 2009584).

### Sequence processing and consolidation into VHVL clones

Individual reads after Guppy base calling typically average more than 10% error per base, and numerous tools exist to align and reduce reads into consensus sequences with substantially improved accuracy^36^. However, such tools were not designed for antibody library sequencing and its huge populations of subtly different sequences which, even assuming successful alignment, group into a myriad of very short, disconnected assemblies. We therefore implemented a bioinformatic pipeline to obtain accurate VHVL sequences from the MinION and iSeq data and to estimate, within each YSD round, the relative abundance of individual VHVL pairs.

Our methods proceed through antibody V(D)J annotation of raw MinION reads using MiXCR (v3.0.13)^37^; iteratively growing and shrinking sequence clusters based on annotated features from each read; sequence error correction and consolidation within each cluster, optionally including high quality Illumina reads; and finally, VHVL clone definition within each sample and quantitation by number of reads mapped to each clone. For enumeration, we only include counts for reads with a length of 1700 to 2100 base pairs. This points to a secondary advantage of MinION sequencing, as shorter reads proliferate during PCR and inflate apparent abundance of particular species. Without length-filtering, relative VHVL abundance calculated from both MinION and iSeq reads are strongly correlated.

### Tissue culture and transient transfection of ECD and RBD

Spike ECD protein and RBD proteins were expressed in Expi293F cells using the manufacturer provided guidelines with slight modifications. In short, a 1 ml frozen working cell bank of Expi293F cells at 1 × 10^7^ viable cells/mL were thawed in a bead bath at 37°C for 2-3 mins. The vial was sprayed with 70% isopropyl alcohol and transferred into a biosafety cabinet. The thawed cells were transferred to a 125 mL non-baffled vented shake flask containing 29 mL of fresh pre-warmed ExpiExpression medium at 37°C. Cells were incubated in at 37°C with ≥80% humidity and 8% CO_2_ on an orbital shaker at 120 rpm and grown until they reached a cell density of 3 × 10^6^ viable cells/ml. Fresh pre-warmed ExpiExpression medium was added to 1 L non-baffled vented shake flask and the 30 mL cell suspension was carefully introduced, making the seeding density of 0.4 × 10^6^ viable cells/mL in a final culture volume of 225 mL. After the cell density reached 3 × 10^6^ viable cells/ml, the culture was expanded to a final volume of 2.25 L in two 2.8 L Thomson Optimum Growth flasks with a seeding density of 0.3 x 10^6^ viable cells/mL. After the cell density reached 3 × 10^6^ viable cells/mL, 2 L of the culture was split into four Thompson flasks with each flask containing 500 mL of culture medium and 500 mL of fresh pre-warmed ExpiExpression medium. The final culture volume in each of the four flasks was 1 L. The cells were incubated at 37°C with ≥80% humidity and 8% CO_2_ on an orbital shaker at 100 rpm. When the cells reached a density of 3 × 10^6^ viable cells/mL, the culture was transferred to two 500 mL sterile centrifuge bottles and the cells were spun down at 100 x g for 10 min. The supernatant was removed, and the cells were resuspended in 4 L of fresh, prewarmed medium. The cells were allowed to re-stabilize in the incubator for 24 h and were transfected at 3 × 10^6^ viable cells/mL in 4 L.

Transfection was performed by diluting 4 mg of plasmid DNA (pDNA) in 240 mL of OptiMEM medium in a sterile bottle and gently inverting 3 - 4 times before incubating at room temperature (RT) for 5 min. ExpiFectamine 293 reagent (13 mL) was then diluted in 225 mL OptiMEM in a sterile bottle and inverted 3 - 4 times before incubating for 5 min at RT. The diluted pDNA and ExpiFectamine reagent were carefully mixed and incubated at RT for 15 min. One-fourth of the combined complex was then slowly transferred to each of the flasks while gently swirling the cells during addition. The cultures were again placed in the incubator with shaking at 100 rpm for 18 h. Post-transfection, ExpiFectamine 293 Transfection Enhancer 1 (6 mL) and ExpiFectamine 293 Transfection Enhancer 2 (60 mL) were added to each flask. The cell viability was monitored every 24 h and the cells were harvested when the viability dropped below 70% or after approximately 3 d. Harvesting was done by centrifugation at 15,900 x g for 45 min at 4°C and the supernatant was transferred into sterile bottles.

### Antigen purification

The feed was prepared adding 0.2 M NaCl and 10 mM imidazole to the supernatant while mixing. The feed material was filtered using a 0.45 µm PES filter membrane pre-wetted with PBS before loading on a prepared Ni-IMAC column. A metal affinity column was prepared by packing IMAC FF beads (Cytiva) into an AxiChrom 70 column housing to a bed height of 9.5 cm and then charging with 50% column volume (CV) of a 0.2 M nickel sulfate solution, washing with water and then 50% CV of 100% “B” buffer (50 mM sodium phosphate buffer containing 300 mM NaCl and 250 mM imidazole, pH 7.8) to remove weakly bound nickel ions. The column was then washed with 100% “A” buffer (50 mM sodium phosphate buffer containing 300 mM NaCl and 20 mM imidazole, pH 7.2) prior to sample loading. The prepared (4.8 L) feed was then loaded onto the column at a linear flow rate of 90 cm/h. After loading, the column was washed with two CVs of 100% A buffer and two CVs of a 13% B buffer (containing 50 mM imidazole) before eluting the protein using 100% B (250 mM imidazole) for 3 CVs. All steps except for the loading step were done at 150 cm/h.

Fractions from the elution peak were pooled and then concentrated 4- to 8-fold by ultrafiltration (UF) using a 115 cm^2^hollow fiber cartridge with either a 50 kDa (S-2P) or a 10 kDa (RBD) molecular weight cutoff membrane (Repligen) and then diafiltered after concentration by exchanging with 5 volumes of PBS.

The concentration of diafiltered protein was determined by measuring the absorbance at 280 nm versus a PBS blank. The protein concentration in mg/mL was obtained using a divisor of 1.03 mL/mg-cm for S-2P and 1.19 mL/mg-cm for RBD. A qualitative assessment of protein quality was made using SDS-PAGE with SYPRO Ruby staining (BioRad) for reduced and non-reduced samples. Only those preparations showing predominantly full-length S-2P (160 kDa subunits) or RBD (70 kDa) were used in ELISAs for assessing neutralizing antibodies.

### Monoclonal antibody expression and purification

VHVL candidates were cloned into custom Golden Gate compatible pCDNA3.4 vectors for IgG1 expression. For transfections, VL was mixed 3:1 with a corresponding VH. Plasmids were transfected into Expi293F (Invitrogen) cells using the recommended protocol. Monoclonal antibodies were harvested at 5-7 days post-transfection. Expi293F cells were centrifuged at 300 x g for 5 min, supernatants were collected and centrifuged at 3000 x g for 20 min at 4℃and diluted to 1X PBS final concentration. Each supernatant was passed through a Protein G or A agarose affinity column (Thermo Scientific). Flow through was collected and passed through the column three times. Columns were washed with 10 CV of PBS and antibodies were eluted with 5 mL 100 mM glycine, pH 2.7 directly in neutralization buffer containing 500 μL 1 M Tris-HCl, pH 8.0.

### Phenotyping assays

Sorted clones from rounds two or three of selection were picked into microplates as described previously. After antibody expression, 10 μL of cells were transferred to a fresh 2-mL microplate and washed 2X with 200 μL cold PBSA buffer. Cells were labeled with 200 nM spike or RBD antigen in 50-100 μL PBSA at RT for 1 h with shaking at 1000 rpm, 3 mm orbit. Labeled cells were washed 2X, and secondary labels were applied as previously described. Cells were resuspended in 200 μL ice-cold PBSA just before analysis. Samples were analyzed on a Sony SA3800 Spectral Cell Analyzer.

### Analysis of mammalian cell surface displayed spike proteins with flow cytometry

The assay has been described in detail in the recent paper by Javanmardi et al^38^. Briefly, plasmids expressing full-length SARS-CoV-2 spike proteins (Spike-Linker-3XFLAG-TM), WT (HexaPro-D614G) and variants, were transfected into HEK293T cells using Lipofectamine 2000 according to manufacturer’s instructions. After 48 h, cells were collected and resuspended in PBS-BSA and incubated with anti-FLAG (mouse) and anti-spike (human) mAbs for 1 h, shaking at RT. Cells were washed 3X with PBS-BSA, resuspended in __ mL and incubated with Alexa Fluor 488 (anti-mouse) and Alexa Fluor 647 (anti-human) antibodies for 30 min, shaking at 4°C. Cells were washed again and resuspended in PBS-BSA prior to flow cytometry analysis (SA3900 Spectral Analyzer, Sony Biotechnology). All data was analyzed with FlowJo (BD Bioscience).

### ELISA

Antigen ELISA plates were made using high-binding plates (Corning, cat. no. 3366) with antigen diluted in PBS to a final concentration of 2 μg/mL. Antigen solution (50 μL) was added to microplates and incubated overnight at 4°C with shaking at 100 rpm, 3 mm orbit. Plates were blocked with PBSM (2% milk in PBS) at RT for 1 h. Plates were washed 3X with 300 μL PBS-T (0.1% Tween-20). Purified antibodies were prepared to 10 μg/mL in PBSM and serially diluted. Antibodies were incubated for 1 h at RT. Plates were washed 3X with PBS-T, and secondary goat anti-human Fab-HRP (Sigma-Aldrich, cat. no. A0293) was applied at 1:5000 in PBSM in 50 μL and incubated at RT for 45 min. HRP substrate (50 μL) was added to wells and the reaction proceeded for 5-15 min until quenched with 50 μL 4M H_2_SO_4_and analyzed for absorbance at 450 nm in a plate reader.

### Live virus neutralization assays

A SARS-CoV-2 microneutralization assay was adapted from an assay used to study Ebola virus^39^. This assay also used SARS-CoV-2 strain WA1. Antibodies were diluted in cell culture medium in triplicate. A SARS-CoV-2 monoclonal antibody was used as a positive control. An antibody that does not bind SARS-CoV-2A was used as negative control. Diluted antibodies were mixed with the SARS-CoV-2 WA1 strain, incubated at 37°C for 1 h, then added to Vero-E6 cells at target MOI of 0.4. Unbound virus was removed after 1 h incubation at 37°C, and culture medium was added. Cells were fixed 24 h post-infection, and the number of infected cells was determined using SARS-CoV-S specific mAb (Sino Biological, cat. no. 401430-R001) and fluorescently-labeled secondary antibody. The percent of infected cells was determined with an Operetta high-content imaging system (PerkinElmer) and Harmonia software. Percent neutralization for each monoclonal antibody at each dilution was determined relative to untreated, virus-only control wells.

Live virus neutralization (VN) for variant of concern B.1.1.7 and WA/1 were performed in an orthogonal assay. The ability of the monoclonal antibodies to neutralize SARS-CoV-2 was determined with a traditional VN assay using SARS-CoV-2 strain USA-WA1/2020 (NR-52281-BEI resources), as previously described^40–43^. All experiments with SARS-CoV-2 were performed in the Eva J Pell BSL-3 laboratory at Penn State and were approved by the Penn State Institutional Biosafety Committee (IBC # 48625). For each mAb a series of 12 two-fold serial dilutions were assessed from a stock concentration of 1 mg/ml. Triplicate wells were used for each antibody dilution. 100 tissue culture infective dose 50 (TCID50) units of SARS-CoV-2 were added to 2-fold dilutions of the diluted mAb. After incubating for 1 hour at 37°C, the virus and mAb mixture was then added to Vero E6 cells (ATCC CRL-1586) in a 96-well microtiter plate and incubated at 37°C. After 3 days, the cells were stained for 1 hour with crystal violet–formaldehyde stain (0.013% crystal violet, 2.5% ethanol, and 10% formaldehyde in 0.01 M PBS). The endpoint of the microneutralization assay was determined as the highest mAb dilution, at which all 3, or 2 of 3, wells are not protected from virus infection. Percent neutralization ability of each dilution of the mAb was calculated based on the number of wells protected, 3, 2, 1, 0 of 3 wells protected was expressed as 100%, 66.6%, 33.3%, or 0%.

### Surface plasmon resonance

To investigate the binding kinetics of mAb N3-1 binding to the spikes, purified His-tagged spike variants (SARS-CoV Tor2 S-2P, SARS-CoV-2 Wuhan-Hu-1 S-HexaPro, SARS-CoV-2 B.1.1.7 S-Hexapro and SARS-CoV-2 B.1.351 S-HexaPro) were immobilized on a Ni-NTA sensor chip (GE Healthcare) using a Biacore X100 (GE Healthcare). For Fab binding experiments, we immobilized spike proteins to a level of ∼450 response units (RUs). Serial dilutions of purified Fab N3-1 were injected at concentrations ranging from 400 to 6.25 nM over spike-immobilized flow cell and the control flow cell in a running buffer composed of 10 mM HEPES pH 8.0, 150 mM NaCl and 0.05% Tween 20 (HBS-T). Between each cycle, the sensor chip was regenerated with 0.35 M EDTA, 50 mM NaOH and followed by 0.5 mM NiCl_2_. For IgG binding experiments, spike immobilization of 200 RUs was used instead to avoid mass transport effect. Serial dilutions of purified IgG N3-1 were injected at concentrations ranging from 25 to 1.56 nM over a spike-immobilized flow cell and the control flow cell. For the SARS-CoV Tor2 S-2P binding experiments, IgG N3-1 concentrations ranging from 100 to 6.25 nM were used. Response curves were double-reference subtracted and fit to a 1:1 binding model or heterogeneous ligand binding model using Biacore X100 Evaluation Software (GE Healthcare).

### Negative stain EM for spike-IgG complexes

To investigate mAb N3-1 binding to spike proteins, purified SARS-CoV-2 Wuhan-Hu-1 S-HexaPro was incubated with 1.2-fold molar excess of IgG N3-1 in 2 mM Tris pH 8.0, 200 mM NaCl and 0.02% NaN_3_ on ice for 10 min. The spike-IgG complexes were at a concentration of 0.05 mg/mL in 2 mM Tris pH 8.0, 200 mM NaCl and 0.02% NaN_3_ prior to deposition on a CF-400-CU grid (Electron Microscopy Sciences) that was plasma cleaned for 30 sec in a Solarus 950 plasma cleaner (Gatan) with a 4:1 ratio of O_2_/H_2_ and stained using methylamine tungstate (Nanoprobes). Grids were imaged at a magnification of 92,000X (corresponding to a calibrated pixel size of 1.63 Å/pix) in a Talos F200C TEM microscope equipped with a Ceta 16M detector. The CTF-estimation, particle picking and 2D classification were all performed in cisTEM^44^.

### Cryo-EM sample preparation and data collection

Purified SARS-CoV-2 S (HexaPro variant) at 0.2 mg/mL was incubated with 5-fold molar excess of Fab N3-1 in 2 mM Tris pH 8.0, 200 mM NaCl and 0.02% NaN_3_ at RT for 30 min. The sample was then deposited on plasma-cleaned UltrAuFoil 1.2/1.3 grids before being blotted for 4 sec with −3 force in a Vitrobot Mark IV and plunge-frozen into liquid ethane. Purified SARS-CoV-2 S with three RBDs covalently trapped in the down conformations (HexaPro-RBD-down variant, S383C/D985C^45–47^) at 0.2 mg/mL, complexed with 2-fold molar excess of Fab A7V3, was deposited on plasma-cleaned UltrAuFoil 1.2/1.3 grids before being blotted for 3 sec with −4 force in a Vitrobot Mark IV and plunge-frozen into liquid ethane. For the HexaPro-N3-1 sample, 3,203 micrographs were collected from a single grid. For the HexaPro-RBD-down-A7V3 sample, 3,636 micrographs were collected from a single grid. FEI Titan Krios equipped with a K3 direct electron detector (Gatan) was used for imaging. Data were collected at a magnification of 22,500x, corresponding to a calibrated pixel size of 1.07 Å/pix. A full description of the data collection parameters can be found in **Extended Data Tables 3-4**.

### Cryo-EM data processing

Gain reference- and motion-corrected micrographs processed by Warp^48^ were imported into cryoSPARC v2.15.0^49^, which was used to perform CTF correction, micrograph curation, particle picking, and particle curation via iterative rounds of 2D classification. The final global reconstructions were then obtained via ab initio reconstruction, iterative rounds of heterogeneous refinement, and subsequently non-uniform homogeneous refinement of final classes with C1 symmetry. For the HexaPro-RBD-down-A7V3 sample, C3 symmetry was attempted in the initial refinement process. Given the low occupancy of the Fabs on a trimeric spike, C1 symmetry was used for the final runs of heterogeneous and homogeneous refinement. To better resolve the Fab-spike interfaces, both datasets were subjected to particle subtraction and focused refinement. Finally, both global and focused maps were sharpened using DeepEMhancer^50^. For A7V3-NTD model building, we used an NTD from the 4A8 complexed spike structure (PDB ID: 7C2L^25^) and a homologous Fab structure (PDB ID: 6IEK) as an initial model to build into map density using UCSF ChimeraX^51^. For N3-1-RBDs model building, we used one RBD-up and one RBD-down from S-HexaPro (PDB ID: 6XKL^52^) and two homologous Fab structures (PDB ID: 5BV7 and 5ITB) as an initial model to build into map density via UCSF ChimeraX. Both models were built further and iteratively refined using a combination of Coot^53^, Phenix^54^, and ISOLDE^55^. The detailed workflows of cryo-EM data processing and data validation can be found in **Extended Data Figures 10, 11 and 14**.

## Ethics statement

The acquisition of blood specimens from convalescent individuals was approved by the University of Texas at Austin Institutional Review Board (protocol 2020-03-085; Breadth of serum antibody immune responses prior to, or following, patient recovery in asymptomatic and non-severe COVID-19). Informed consent was obtained from all participants.

## Data availability

The sequences of neutralizing antibodies have been deposited in GenBank with accession numbers. Molecular coordinates for A7V3 and N3-1 Fab complexes with SARS-CoV-2 trimeric spike protein have been deposited to the Protein Data Bank. Structural data are presented in Figs. 4-5, Extended Data Tables 3-4, and Extended Data Figs. 10-12 and 14.

## Author Contributions

Conceptualization: JG, CH, AH, ECG, DRB, JSM, and JDG; Methodology: JG, CH, AH, ECG, FB, NW, KJ, AH, RR, MJJ, SLW, ZLN, JL, TSS, SVK, VK, RAH, IF, DRB, JSM, and JDG; Investigation: JG, CH, AH, ECG, FB, NW, AA, KJ, AH, WNV, JAC, ACM, RR, MJJ, SLW, ZLN, JL, RAH, SVK, VK, IJF, JMM, DRB, JSM, and JDG; Data Analysis and Interpretation: JG, CH, AH, DRB, NW, AH, JMD, IJF, EMM, JSM, and JDG; Data Curation: JG, CH, AH, ECG, NW, DRB, JSM, AH, JDM, JSM, and JDG; Original Draft: JG, CH, AH, ECG, DRB, JMM, JSM, VK, EMM, and JDG; Review & Editing: JG, CH, AH, ECG, DRB, ADE, EMM, JMM, IJF, GG, JSM, and JDG; Funding: JMD, IJF, GCI, VK, SVK, GG, ADE, JSM and JDG.

## Acknowledgements

We would like to thank the many frontline responders, who have sacrificed tirelessly for the health and treatment of COVID-19 patients. In particular, we would like to extend our gratitude to Daniel Billick for acquiring blood samples from each of the patients represented here. We thank Dr. Kathryn Stockbauer for careful editing of the manuscript. We thank Samantha Nagel for her tremendous efforts. We also would like to thank Dr. Sasha Dickinson from the Sauer Structural Biology Laboratory at the University of Texas at Austin for his assistance with microscope data collection. The Sauer Structural Biology Laboratory is supported by the University of Texas College of Natural Sciences and by award RR160023 from the Cancer Prevention and Research Institute of Texas (CPRIT). Funding for USAMRIID was provided through the CARES Act with programmatic oversight from the Military Infectious Diseases Research Program–project 14066041. Opinions, conclusions, interpretations, and recommendations are those of the authors and are not necessarily endorsed by the U.S. Army. The mention of trade names or commercial products does not constitute endorsement or recommendation for use by the Department of the Army or the Department of Defense. Molecular graphics and analyses were performed with UCSF Chimera, developed by the Resource for Biocomputing, Visualization, and Informatics at the University of California, San Francisco, with support from NIH P41-GM103311. This research was funded in part by: the Army Research Laboratory’s TRANSFORME Essential Research Program (JDG, DRB, RR, THSS); a Cooperative Agreement (W911NF-17-2-0091) between ARL and UT Austin to ADE, EMM, and GG; a grant from DTRA (HDTRA12010011) awarded to JDG and ADE; a National Institutes of Health (NIH)/National Institute of Allergy and Infectious Diseases (NIAID) grant awarded to JSM (R01 AI127521); financial assistance award 70NANB20H037 to ZLN, SLW, and JDG from the US Department of Commerce, National Institutes of Standards and Technology (NIST) via the National Institute for Innovation in Manufacturing Biopharmaceuticals (NIIMBL); a Welch grant (F-1016) awarded to IJF; a Welch grant (F-1515) awarded to EMM; funding support from the Huck Institutes of Life Sciences and the Pennsylvania Agricultural Experiment Station (to VK and SK); an NIH grant (R01 AI158177-01) to SK; and an NIH grant (R35 GM122480) to EMM. GCI expended discretionary funds from his MBS-20 account.

