## Supplementary Information for "Synthetic repertoires derived from convalescent COVID-19 patients enable discovery of SARS-CoV-2 neutralizing antibodies and a novel quaternary binding modality"

**Extended Data**

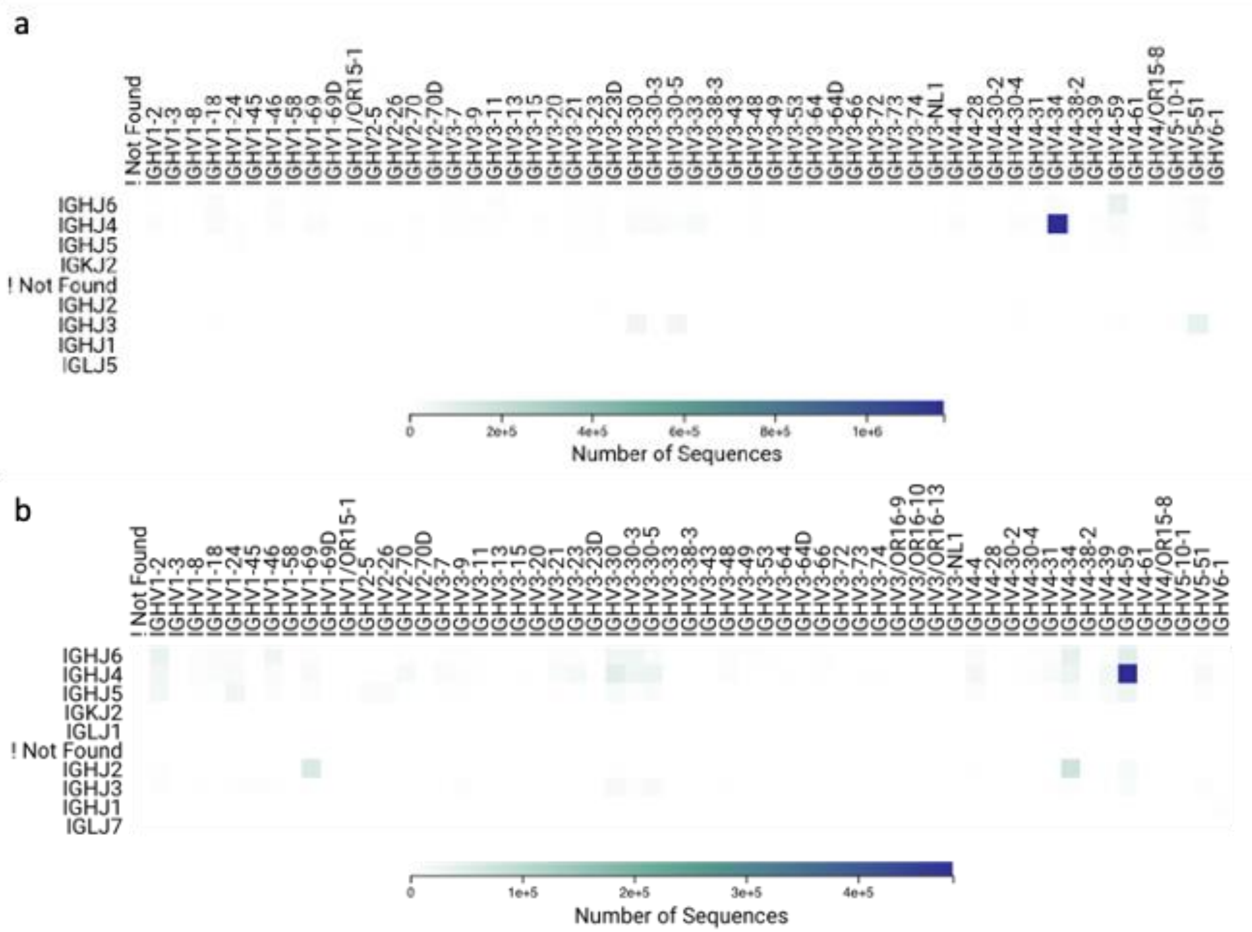


**Extended Data Figure 1. Summary of IgG VH-only sequencing of donor 1 [a] and donor 2 [b].** Libraries generated from these sequences were used for IgSeq proteomics and as a starting point for YSD selections.

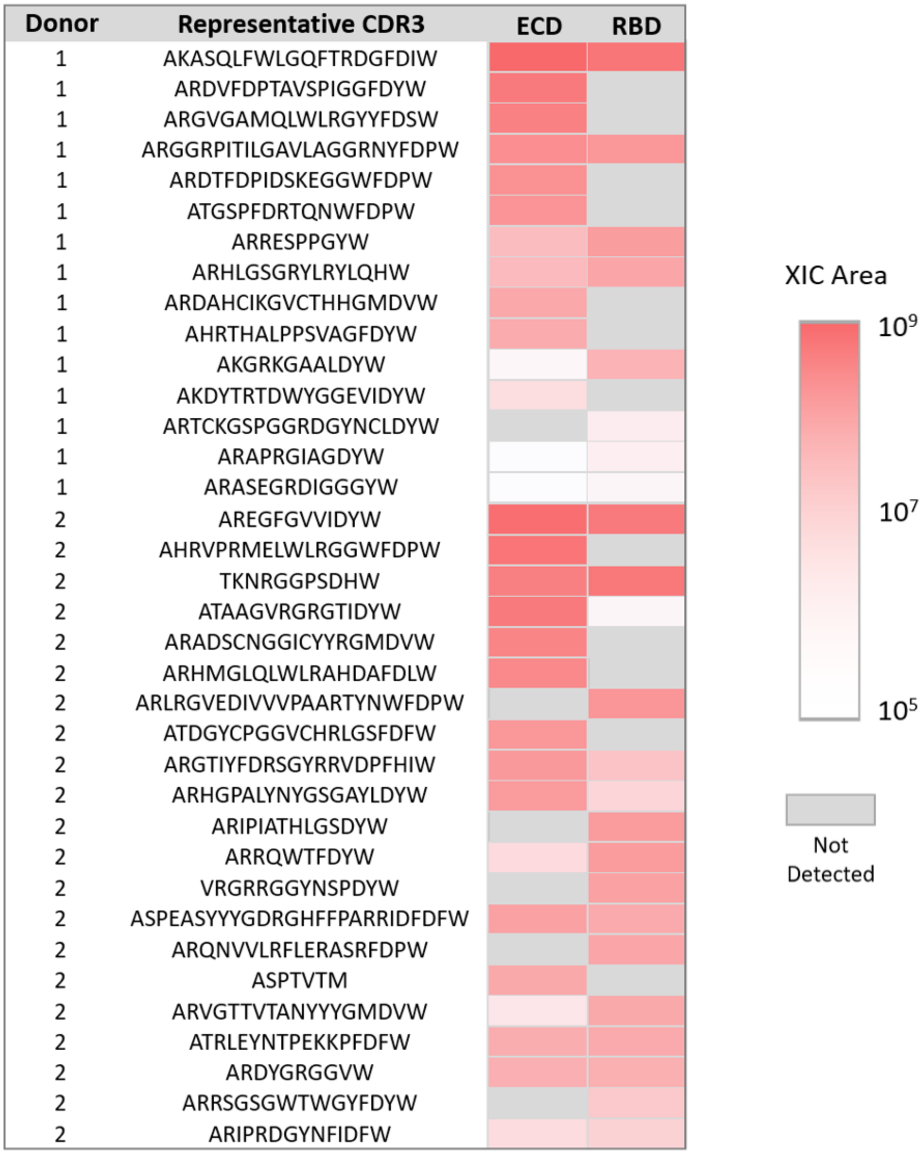


**Extended Data Figure 2. Anti-SARS-CoV-2 ECD and RBD antibody clonotypes identified in the serum of patients 1 and 2 (P1, P2) by Ig-Seq proteomic analysis.** Heat maps represent the relative abundances of unique clonotypes calculated as the sum of XIC peak areas of CDR3-peptides observed by LC-MS/MS.

| Name | V-Gene | J-Gene | Light CDR1 | Light CDR2 | Light CDR3 |
|---|---|---|---|---|---|
| PLC1 | IGKV1-5 | IGKJ1 | QSISSW | DAS | QQYNSYSPWT |
| PLC2 | IGKV2-28 | IGKJ2 | QSLLHSNGYNY | LGS | MQALQTPPYT |
| PLC3 | IGKV3-11 | IGKJ4 | QSVSSY | DAS | QQRSNWPPLT |
| PLC4 | IGKV3-15 | IGKJ1 | QSVSSN | GAS | QQYNNWPPWT |
| PLC5 | IGKV3-20 | IGKJ1 | QSVSSSY | GAS | QQYGSSPPWT |
| PLC6 | IGKV4-1 | IGKJ2 | QSVLYSSNNKNY | WAS | QQYYSTPPYT |
| PLC7 | IGLV1-44 | IGLJ3 | SSNIGSNT | SNN | AAWDDSLNGPVV |
| PLC8 | IGLV1-51 | IGLJ3 | SSNIGNNY | DNN | GTWDSSLSAVV |
| PLC9 | IGLV3-1 | IGLJ3 | KLGDKY | QDS | QAWDSSTVV |

**Extended Data Table 1. List of public light chains screened in this study.** Light chain V-genes were derived from a previously published dataset of homeostatic repertoires from three donors. For each of these V-genes, its most commonly associated germline J-gene was chosen.

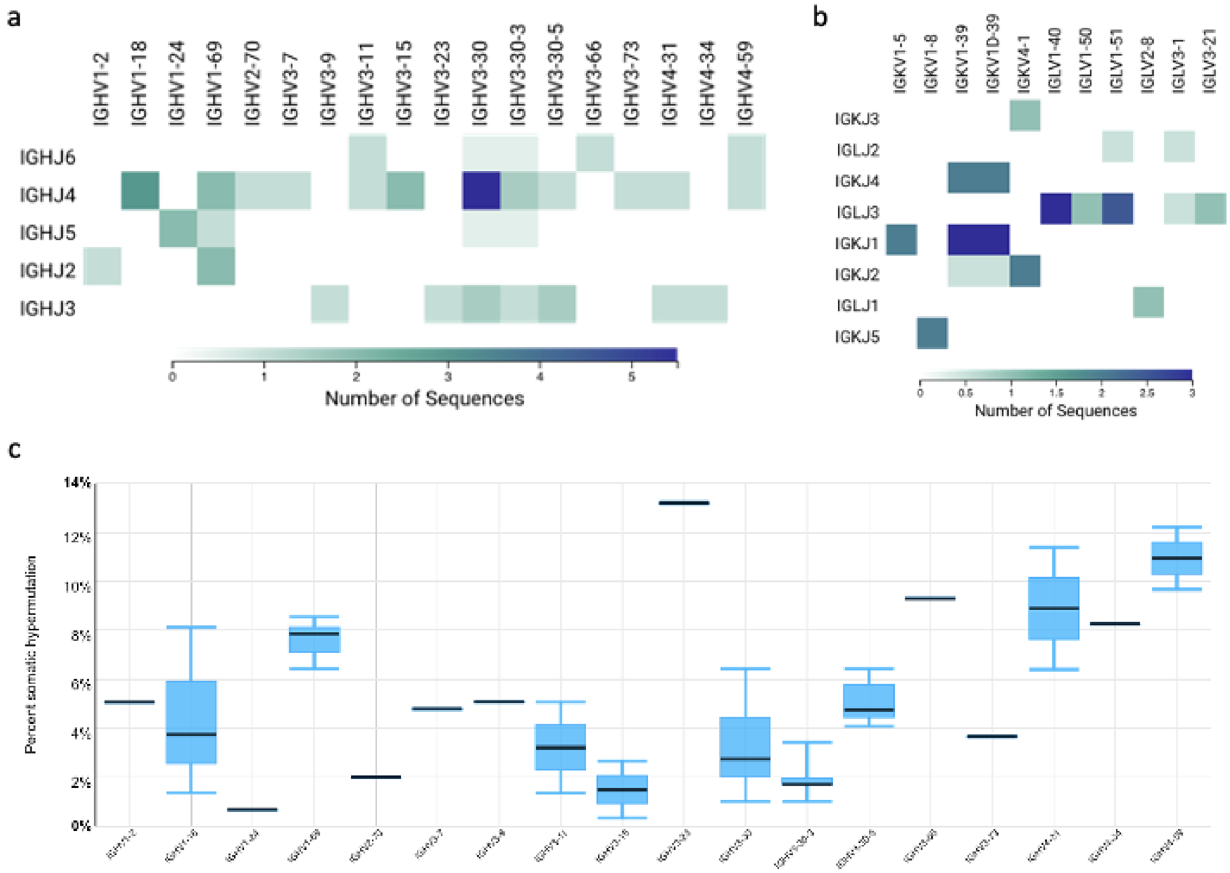


**Extended Data Figure 3. a,** IgSeq VH gene usage. **b,** YSD-IgSeq VL gene usage. **c,** Somatic hypermutation as a function of gene family calculated using Geneious Biologics.

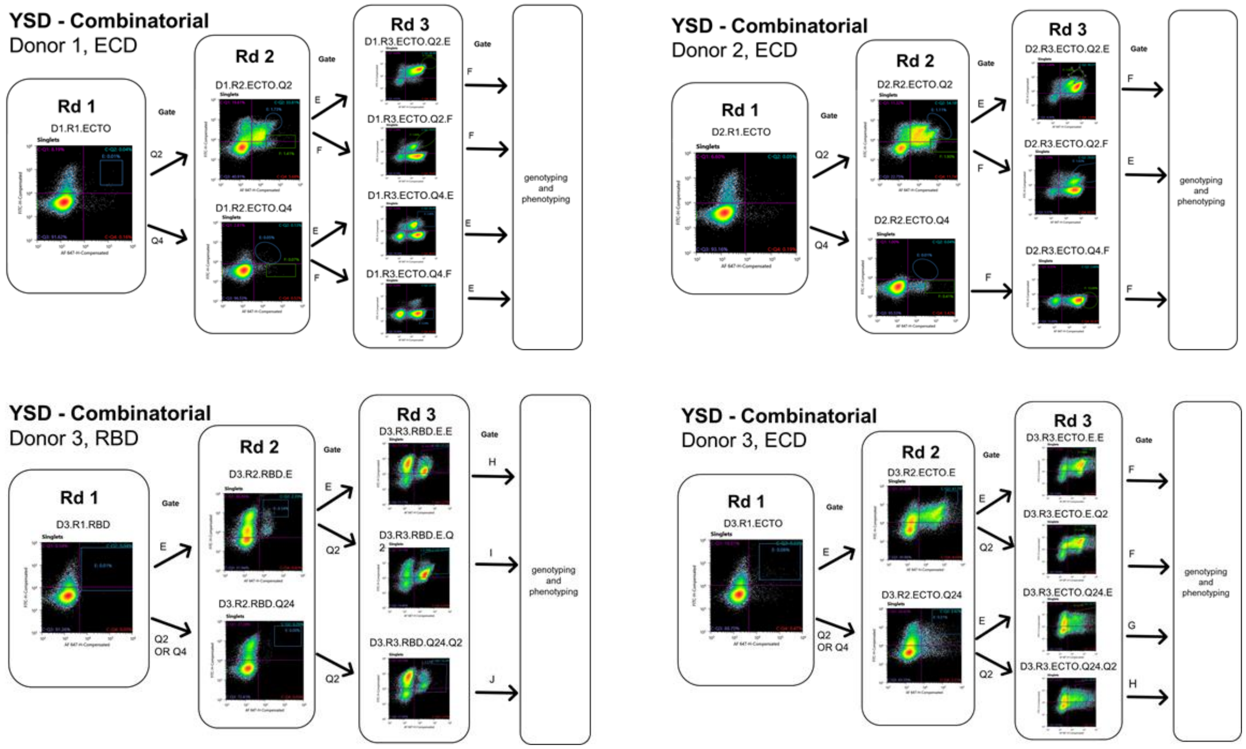


**Extended Data Figure 4. Representative YSD cell sorting lineage plots**. Donor repertoires were cloned as Fab libraries and displayed in yeast. Yeast were labeled for expression (y-axis, anti-FLAG-FITC) and antigen binding (x-axis, biotinylated ECD, Streptavidin-Alexa Flour 488; human-Fc RBD, anti-human Alexa Fluor 488). Each library was subjected to selection in the presence of either RBD or spike ECD. Each population was subjected to sorting into up to two gates at a time, such that after several rounds of selection, populations with various expression and binding characteristics were enriched (see Rd 3). Combinatorial libraries consisted of randomly-paired VHs and VLs cloned from donor cDNA.

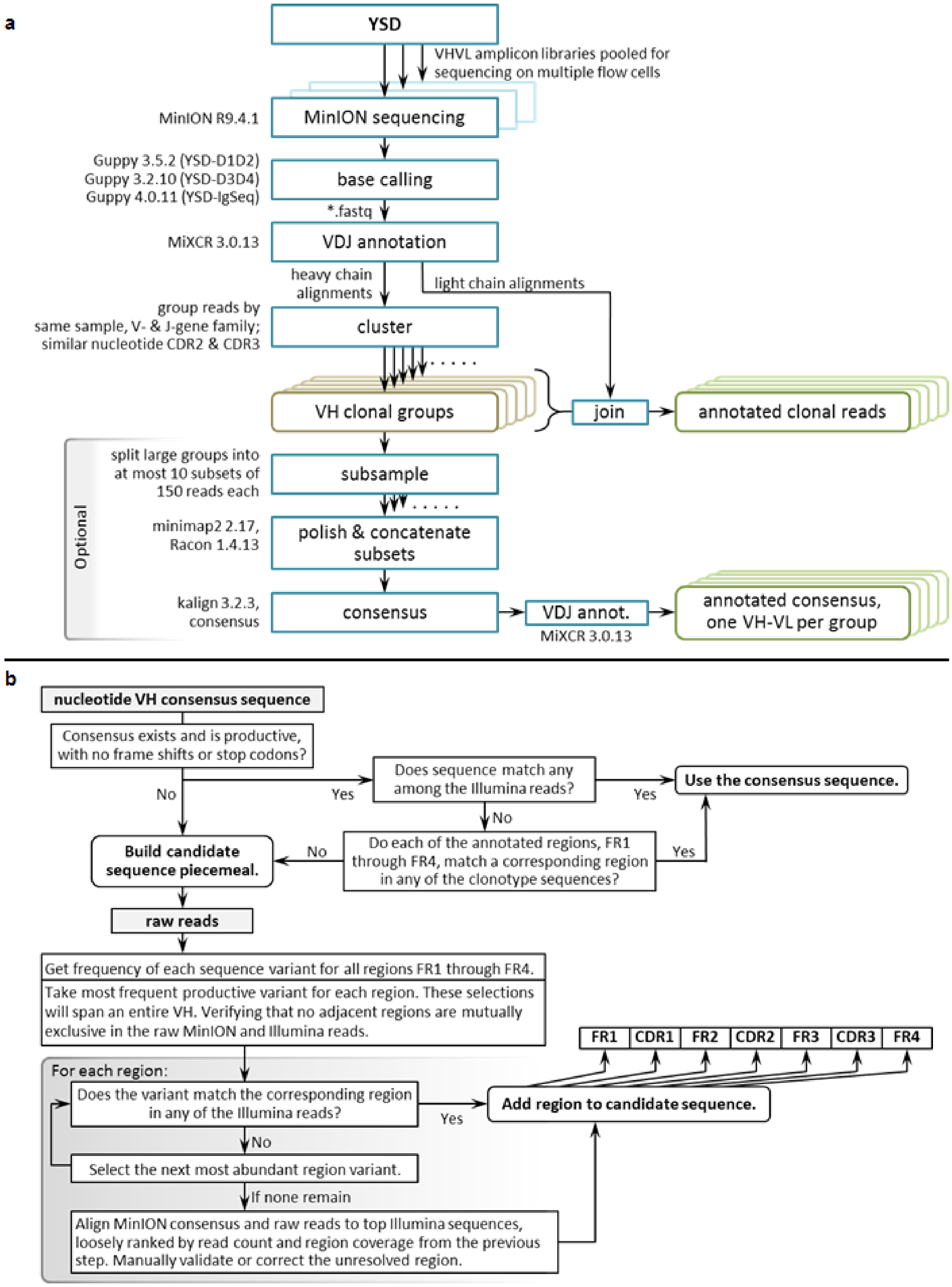


**Extended Data Figure 5. Definition of candidate VH-VL sequences from error-prone MinION sequencing.**

| VH CDR3 | MinION 1.7-2.1kb total reads | MinION all total reads | iSeq total reads |
|---|---|---|---|
| AKTSGYNLPDY | 2467936 | 3218688 | 82347 |
| ARGTIYFDRSGYRRVDPFHI | 682539 | 1153588 | 41256 |
| ATGPAVRRGSWFDP | 189383 | 250994 | 14595 |
| ARRQWTFDY | 170557 | 198948 | 5961 |
| ATGSPFDRTQNWFDP | 150096 | 195000 | 9471 |
| ARDGGGYVSY | 137057 | 184361 | 7760 |
| ARGVGAMQLWLRGYYFDS | 101311 | 136418 | 10220 |
| TTDRTYDYVWGSYRYEDY | 18251 | 26192 | 1497 |
| ASPEASYYYGDRGHFFPARRIDFDF | 14123 | 19357 | 1356 |
| TTDRSYDYVWGSYRYPDY | 13957 | 20327 | 1046 |
| ARIPIATHLGSDY | 13507 | 20181 | 1058 |
| ATAAAVRGRGTIDY | 11135 | 17518 | 924 |
| ARDYGRGGV | 9303 | 12940 | 631 |
| AKDRAIFYLNPRYYLDY | 9245 | 12123 | 881 |
| ASPTVT | 8847 | 11590 | 436 |
| ARELPPGRMVVPATYWHFDL | 7591 | 11084 | 928 |
| TSPRADYGSNSVGNY | 5736 | 7607 | 725 |
| ARDGITNSGVVTHFGMDV | 4214 | 5713 | 320 |
| ARHMGMQLWLRAHDAFDL | 908 | 1137433 | 46761 |
| ARLGTGYP | 655 | 1077 | 32 |
| ACSRARGFMFFDY | 631 | 1850171 | 71835 |
| AKATQLFWLGQFTRDGFDI | 366 | 2110 | 128 |
| ARADSCNGGICYYQGMDV | 215 | 602990 | 27489 |
| AREQPPGRMVVPATYWHFDL | 70 | 8668 | 222 |

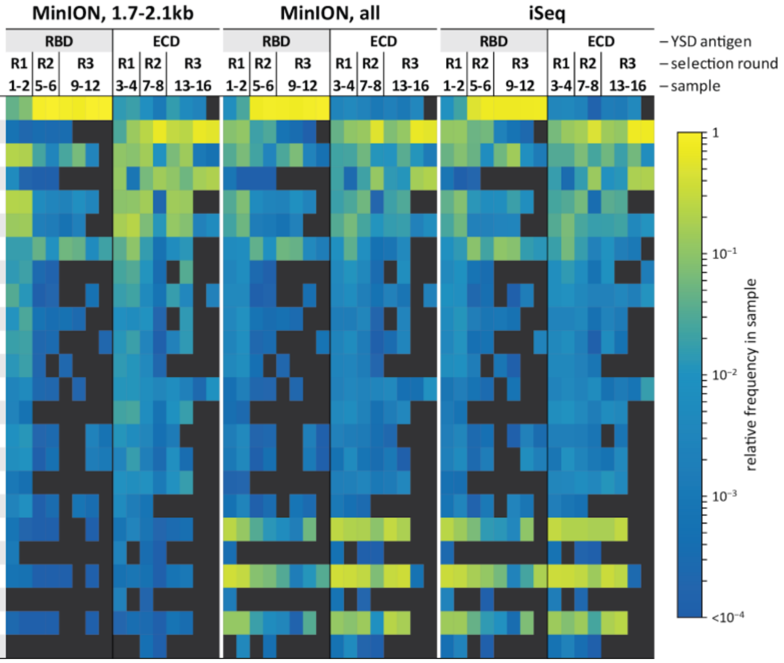


**Extended Data Figure 6. VH CDR3 read counts and relative abundance from the YSD-IgSeq experiment**, shown for MinION reads of 1.7 kb to 2.1 kb (left), all MinION reads (center) and all iSeq reads (right). Heatmap values are CDR3 frequencies, or read counts normalized within each respective sample. Length requirements improve VHVL abundance estimation by, in particular, removing the inflated counts of short artifactual reads due to unbalanced PCR amplification. Comparison with and without filtering shows certain CDR3 are disproportionately affected. The iSeq read counts for a VH CDR3 include only reads with identical amino acid CDR3 annotation. The respective MinION counts also include reads diversified through sequencing error, recovered through our clustering methods. High MinION error rates create enormous sequence diversity. Full-length VH diversity is bounded only by the total read count. (center, right panels) There is strong correspondence between the unfiltered MinION and iSeq CDR3 frequencies. This indicates that our independent MinION read consolidation techniques successfully recapitulate CDR3 patterns observed from the much more accurate Illumina sequencing platform.

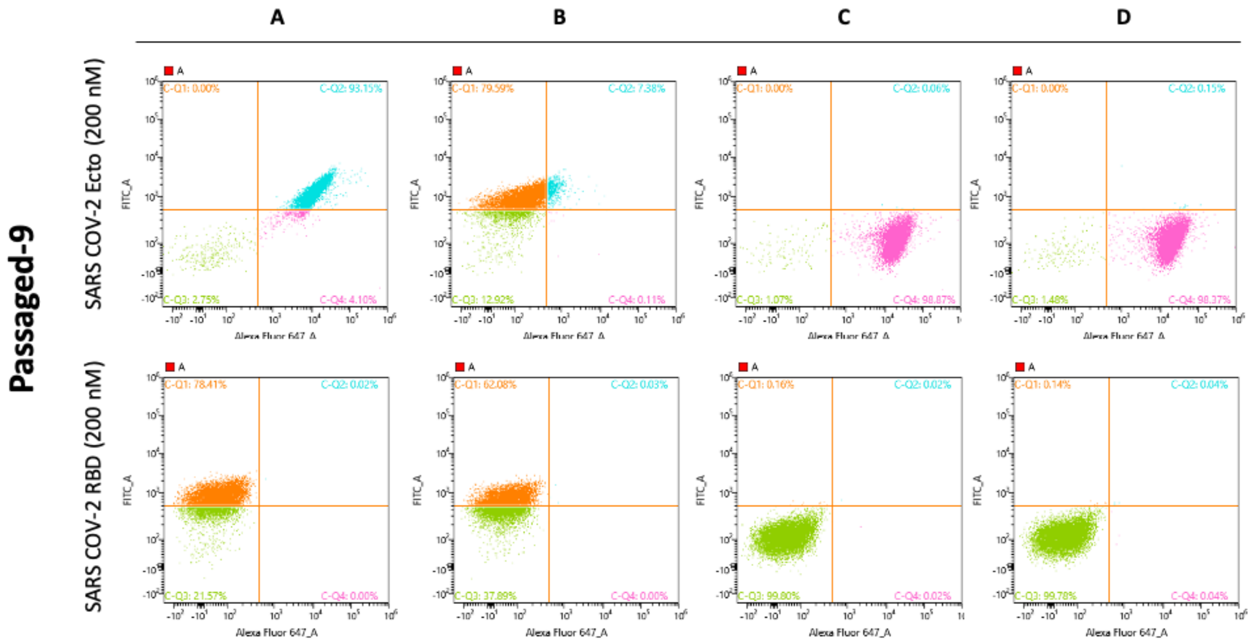


**Extended Data Figure 7. Representative YSD single clone flow cytometry.** The x-axis is binding, and the y-axis is expression. Each of the four columns represents a single clone assayed against Ecto (top row) and RBD (bottom row). Sequencing confirmed clones **c** and **d** are identical. Clones were initially prioritized on expression normalized binding and lack of RBD binding.

a

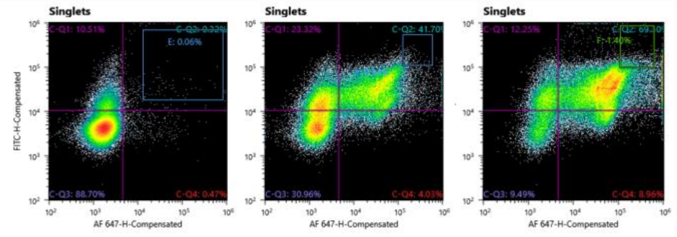


b

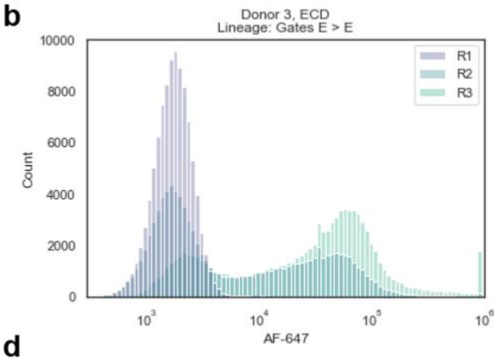


c

| Rank | CDRH3 | Enrichment factor | Rank by read count | Change |
|---|---|---|---|---|
| 1 | ATASPHCSSTSCSINWFDP | 9988 | 2 | 1 |
| 2 | ASGGGSYYSPFDY | 2413 | 7 | 5 |
| 3 | ARAGGGGYQGPFDY | 1991 | 1 | -2 |
| 4 | ARAYGGGYLTPFDY | 1391 | 4 | 0 |
| 5 | AKPQGGGYNGAFDI | 720 | 6 | 1 |
| 6 | ATVSPFWSGYYNNWFDP | 407 | 3 | -3 |
| 7 | ARPVTGGYYYYMDV | 380 | 8 | 1 |
| 8 | TTERYYDYIWGSYRYSDY | 342 | 10 | 2 |
| 9 | ARDLRGSYSGLLDY | 307 | 11 | 2 |
| 10 | **ATRFAVYGDYLIDY** | 286 | 9 | -1 |
| 11 | AKVVRHLYSSGYSNWFDP | 233 | 20 | 9 |
| 12 | ANQVTYYDYIWGSYRPGAFDI | 150 | 13 | 1 |
| 13 | ARDKGRGILFTHRDAFDI | 140 | 29 | 16 |
| 14 | ARRRVYPSGFDY | 138 | 18 | 4 |
| 15 | ARASHGGYSSAFDP | 131 | 15 | 0 |
| 16 | VRSGHCNGGVCYGAEYFQH | 112 | 31 | 15 |
| 17 | ARDGSGGYSNPDY | 111 | 12 | -5 |
| 18 | VKGAGGYFDS | 110 | 32 | 14 |
| 19 | ATAPAVVVGNWFDP | 94 | 34 | 15 |
| 20 | ARGLDIIVVPAALDH | 92 | 5 | -15 |

d

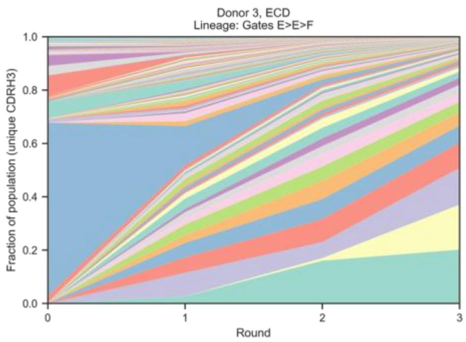


**Extended Data Figure 8. Yeast surface display of a combinatorially assembled donor repertoire.**

**a,** Representative example of YSD cell sorting showing donor 3 Fab selection with the ECD antigen. The x-axis (AF-647) shows antigen binding and the y-axis (FITC) shows Fab expression level. Each round was sorted using one or two gates to enrich populations with different phenotypes. **b**, Histogram showing enrichment of binders from round one (purple) to round 3 (light green) in the highlighted lineage. **c**, Table showing the top HCDR3s as ranked by enrichment in this lineage. Relative ranking by raw readcount is also compared. The bolded HCDR3 represents neutralizing mAb 7-6. **d**, Area chart showing HCDR3 enrichment throughout selection. Each unique HCDR3 is represented as a fraction of all HCDR3s in that population. Significant bias exists in the initial library, but by the end of the selection top variants represent over 10% of the total population.

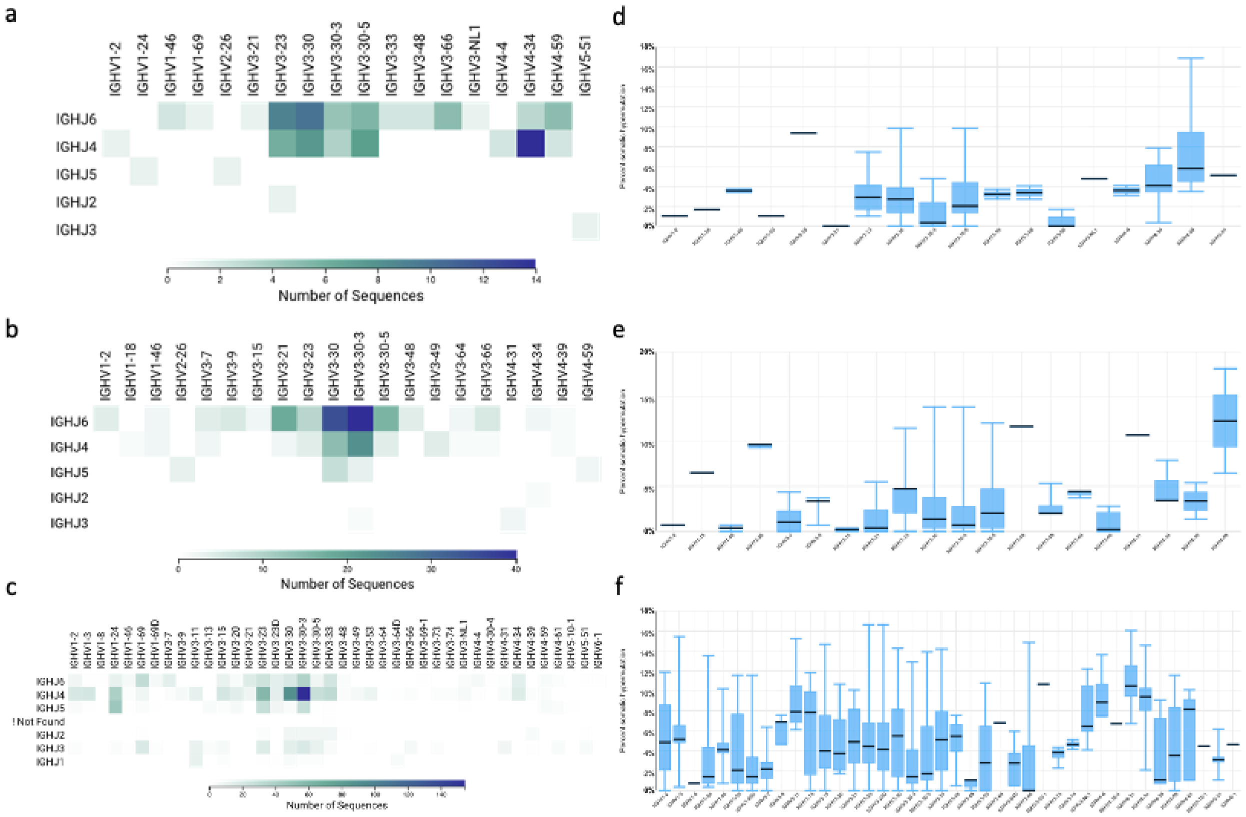


**Extended Figure 9. a-c,** YSD selected VHs from donors 1-3 after three rounds of selection. **d-f,** Somatic hypermutation by gene family as calculated by Geneious Biologics.

**Extended Data Table 2. Summary of neutralizing antibodies**

| mAb | IC50 nM | VH gene | H-CDR3 | VL gene | L-CDR3 | Epitope | HC source | LC source |
|---|---|---|---|---|---|---|---|---|
| **8-32** | 0.018 | IGHV1-24 | ATGSPFDRTQNWFDP | IGLV2-8 | SSYAGSNNLA | NTD | IgSeq | YSD |
| **N3-1** | 0.25 | IGHV4-31 | ARGTIYFDRSGYRRVDPFHI | IGKV1-5 | QQYNSYSPWT | RBD | YSD | PLC |
| **8-131** | 0.30 | IGHV1-24 | ATGSPFDRTQNWFDP | IGLV1-51 | GTWDNSLSAGV | NTD | IgSeq | YSD |
| **8-114** | 0.35 | IGHV1-24 | ATGPAVRRGSWFDP | IGLV1-51 | GTWDSSLSGYV | NTD | IgSeq | YSD |
| **12C8** | 0.84 | IGHV1-24 | ATGPAVRRGSWFDP | IGLV1-51 | GTWDSSLSAVV | NTD | IgSeq | PLC |
| **A7V3** | 0.95 | IGHV1-24 | ATGSPFDRTQNWFDP | IGLV1-51 | GTWDSSLSAVV | NTD | IgSeq | YSD |
| **7-6** | 0.98 | IGHV1-24 | ATRFAVYGDYLIDY | IGLV3-19 | NSRDSSGDLVV | NTD | YSD | YSD |
| **8-132** | 1.39 | IGHV1-24 | ATGSPFDRTQNWFDP | IGLV1-51 | GTWDSSLSAGV | NTD | IgSeq | YSD |
| **4C7** | 2.15 | IGHV1-24 | ATAAAVRGRGTIDY | IGLV1-44 | AAWDDSLNGPVV | NTD | IgSeq | PLC |
| **4C8** | 2.35 | IGHV1-24 | ATAAAVRGRGTIDY | IGLV1-51 | GTWDSSLSAVV | NTD | IgSeq | YSD |
| **7A8** | 4.24 | IGHV1-24 | ATGSPFDRTQNWFDP | IGLV1-51 | GTWDSSLSAVV | NTD | IgSeq | PLC |
| **8-3** | 6.42 | IGHV3-66 | ARGGVVDTYYYYGMDV | IGLV7-46 | LLSQSGAWV | NTD | IgSeq | YSD |
| **3-26** | 9.42 | IGHV3-30-3 | ARPYSGSYWGYFDY | IGLV1-47 | AAWDDSLSGPV | S2 | YSD | YSD |
| **N3-3** | 13.1 | IGHV4-31 | ARGTIYFDRSGYRRVDPFHI | IGKV3-11 | QQRSNWPPLT | RBD | YSD | PLC |
| **1D4** | 25.5 | IGHV2-70 | ARIPIATHLGSDY | IGKV3-15 | QQYNNWPPWT | RBD | IgSeq | PLC |
| **N3-7** | 27.7 | IGHV4-31 | ARGTIYFDRSGYRRVDPFHI | IGLV1-44 | AAWDDSLNGPVV | RBD | YSD | PLC |
| **6-3A** | 32.0 | IGHV4-31 | ARGTIYFDRSGYRRVDPFHI | IGLV8-61 | TLYMGGGLLV | RBD | YSD | YSD |
| **8-96** | 33.5 | IGHV3-30 | AKAPGQWLRFHYYGMDV | IGLV1-40 | NSRDINSNHVL | NTD | IgSeq | YSD |
| **8B5** | 34.8 | IGHV1-2 | ARELPPGRMVVPATYWHFDL | IGKV3-20 | QQYGSSPPWT | RBD | IgSeq | PLC |
| **3B9** | 51.4 | IGHV3-30 | ARDGGGYVSY | IGLV3-1 | QAWDSSTVV | NTD | IgSeq | PLC |
| **8-130** | 65.1 | IGHV3-30 | ATGPAVRRGSWFDP | IGKV1-39 | QQSYSTPRT | NTD | IgSeq | YSD |
| **4A5** | 65.7 | IGHV3-30-3 | AKASQLFWLGQFTRDGFDI | IGKV3-20 | QQYGSSPPWT | RBD | IgSeq | PLC |
| **6-3B** | 81.5 | IGHV4-31 | ARGTIYFDRSGYRRVDPFHI | IGLV1-40 | QSYDGSLNDDVI | RBD | YSD | YSD |
| **B3.1** | 124 | IGHV3-30 | ARARGGSYYYGMDV | IGKV1-5 | QQYNSYSPWT | S2 | YSD | PLC |
| **8-42** | 124 | IGHV3-30 | ARDYGRGGV | IGKV1-39 | QQSYSTRPLT | NTD | IgSeq | YSD |
| **1D1** | 173 | IGHV2-70 | ARIPIATHLGSDY | IGKV1-5 | QQYNSYSPWT | RBD | IgSeq | PLC |
| **1D9** | 183 | IGHV2-70 | ARIPIATHLGSDY | IGLV3-1 | QAWDSSTVV | RBD | IgSeq | PLC |
| **N6-2** | 222 | IGHV3-30-3 | ARPYSGSYWGYFDY | IGKV2-28 | MQALQTPPYT | S2 | YSD | PLC |
| **P4D3** | 242 | IGHV3-30 | AKAPGQWLRFHYYGMDV | IGLV1-40 | QSYGNNQGV | S2 | YSD | YSD |
| **P4A3** | 272 | IGHV3-30 | ARDDTGRVGSGWYCPLY | IGKV1-39 | QQSYSTPLS | S2 | YSD | YSD |
| **8-19** | >300 | IGHV4-31 | ARGTIYFDRSGYRRVDPFHI | IGKV1-39 | QQSYSTRPLT | RBD | YSD | YSD |
| **3-18** | >300 | IGHV3-30 | AKQAGAYCSGGSCYSSSEADY | IGLV1-40 | QSYGNNQGV | RBD | YSD | YSD |
| **P3B10** | >300 | IGHV3-30-3 | ARPYSGSYWGYFDY | IGLV3-1 | QAWDSSTF | S2 | YSD | YSD |
| **6-3C** | >300 | IGHV4-31 | ARGTIYFDRSGYRRVDPFHI | IGLV3-1 | QAWDSSTVV | RBD | YSD | YSD |
| **4A7** | >300 | IGHV3-30-3 | AKASQLFWLGQFTRDGFDI | IGLV1-44 | AAWDDSLNGPVV | RBD | IgSeq | PLC |
| **1D5** | >300 | IGHV2-70 | ARIPIATHLGSDY | IGKV3-20 | QQYGSSPPWT | RBD | IgSeq | PLC |
| **P3C6** | >300 | IGHV3-23 | APGRSLY | IGLV1-40 | QSYDSGLSGSI | S2 | YSD | YSD |

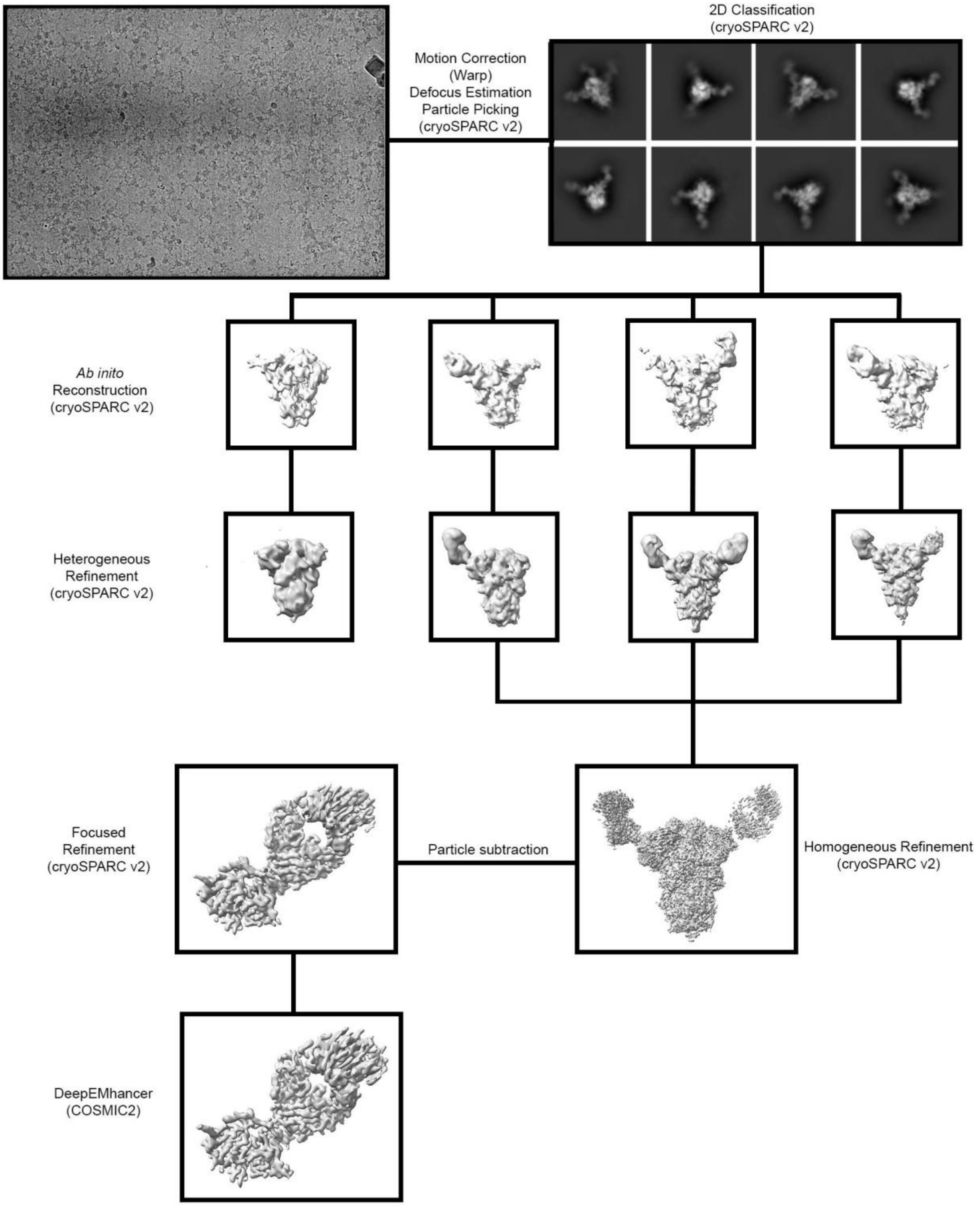


**Extended Data Figure 10**. **Cryo-EM data processing workflow for A7V3 bound SARS-CoV-2 S.**

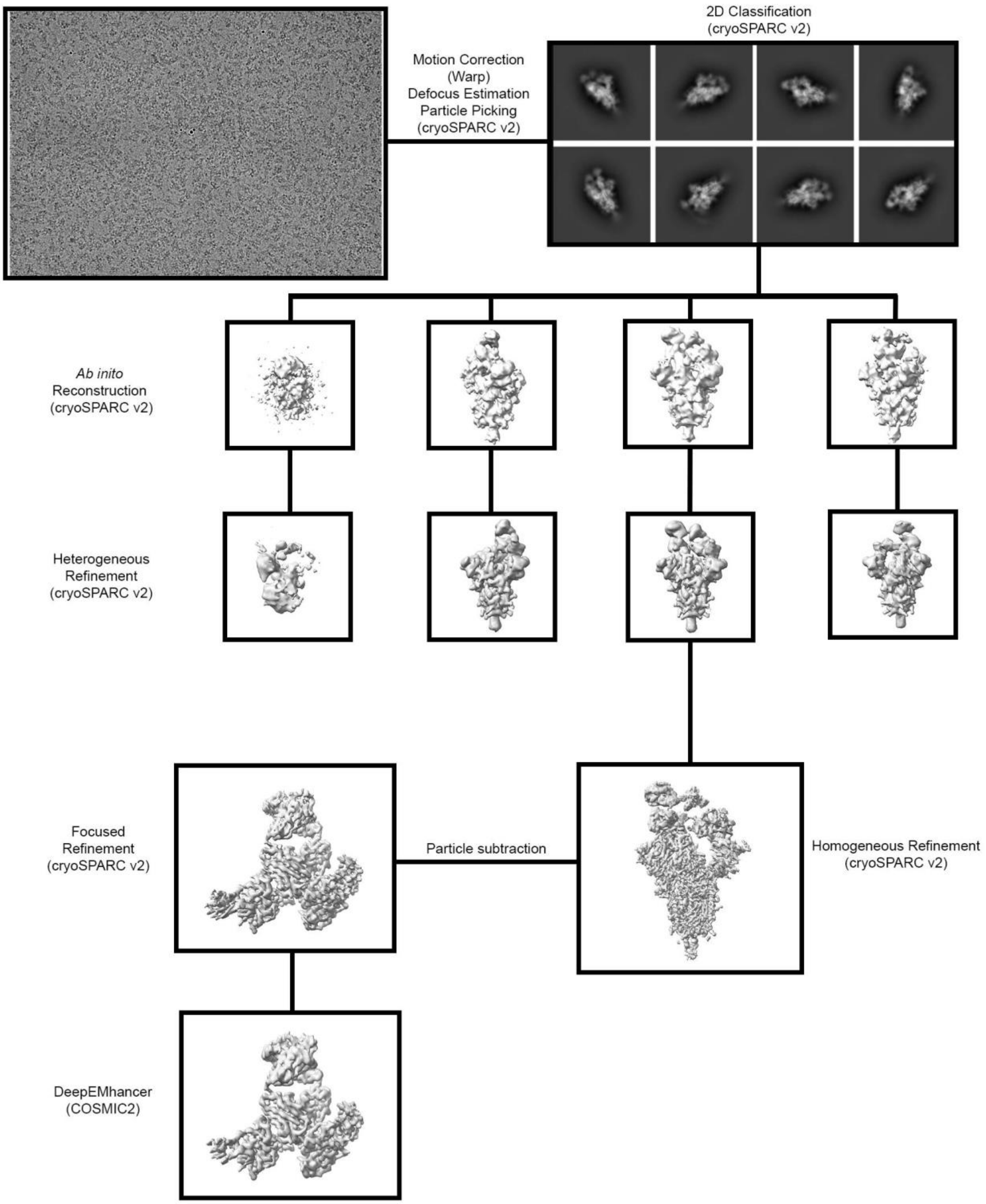


**Extended Data Figure 11**. **Cryo-EM data processing workflow for N3-1 bound SARS-CoV-2 S.**

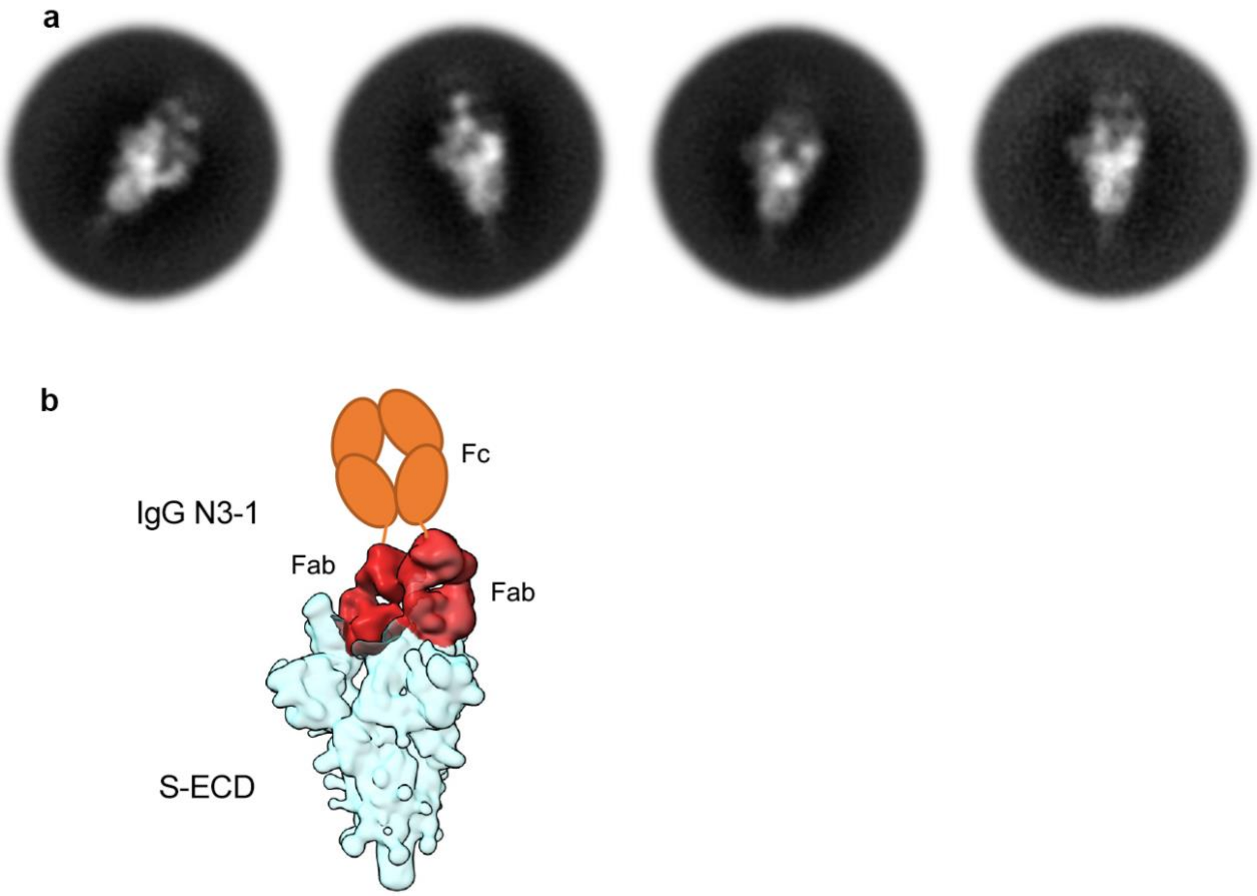


**Extended Data Figure 12**. **Avidity of mAb N3-1 likely achieved by a single IgG binding to a trimeric spike**. **a,** Representative 2D class averages of IgG N3-1 complexed with SARS-CoV-2 S by negative stain electron microscopy (nsEM). Although the density of Fc is not well-resolved, two clear densities of Fabs are clearly visible per trimeric spike. **b,** A schematic model generated by Gaussian-smoothened cryoEM map of N3-1 bound to SARS-CoV-2 S. The Fab density is highlighted in brick red, and the spike is shown in light blue. The unobserved Fc is shown as orange ovals.

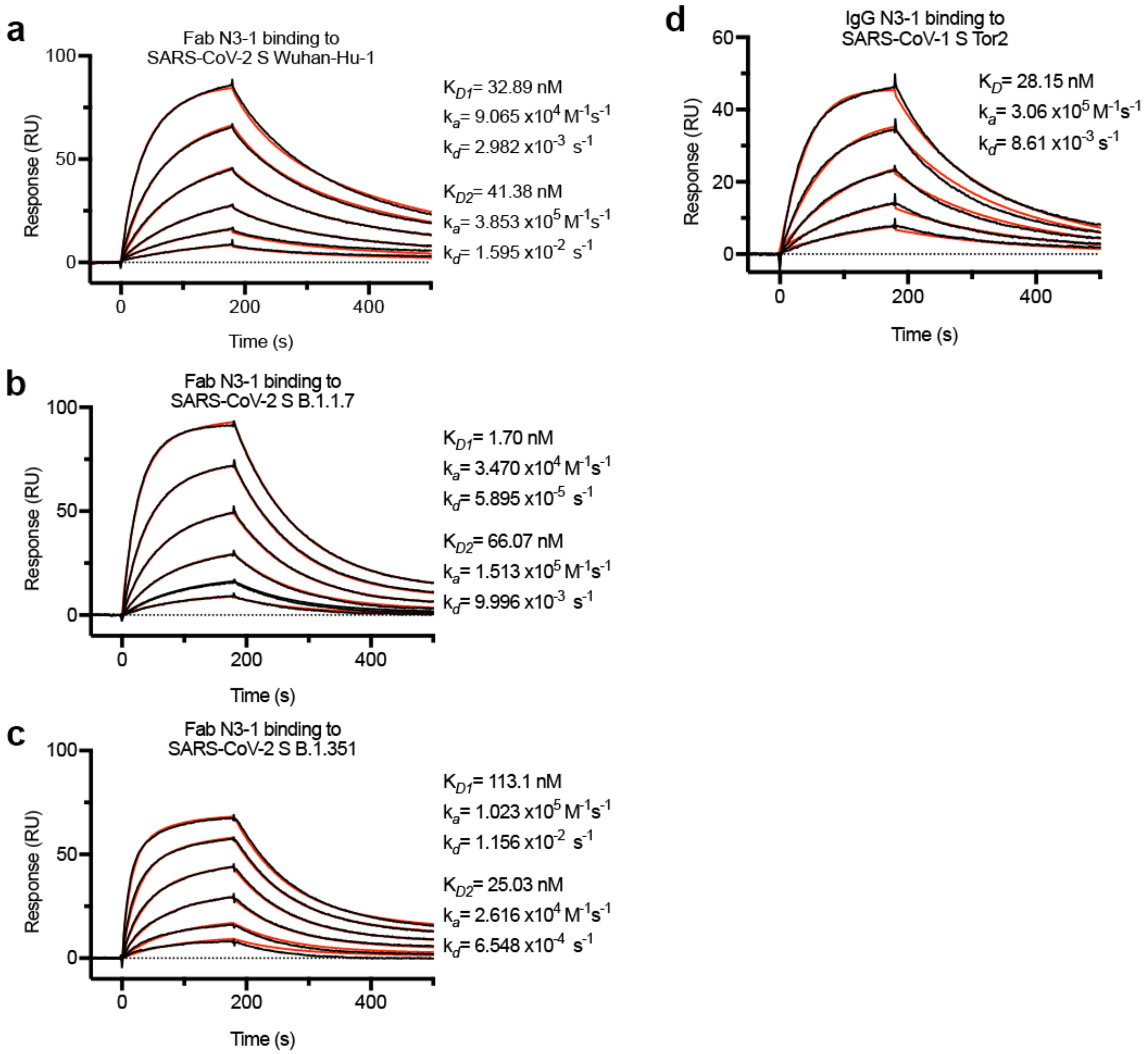


**Extended Data Figure 13**. **mAb N3-1 exhibits cross-reactivity and avidity to CoV spikes**
**a-c,** Binding of Fab N3-1 to SARS-CoV-2 S WuHan-Hu-1 **[a]**, its variants B.1.1.7 **[b]** and B.1.351 **[c]** were assessed by surface plasmon resonance (SPR) using an NTA sensor chip. **d,** Binding of IgG N3-1 to SARS-CoV-1 S was also assessed by SPR. Binding data are shown as black lines. For **[a-c]**, the best fit to a heterogeneous binding model is shown as red lines. For **[d]**, the best fit was achieved using a 1:1 binding model and shown as red lines.

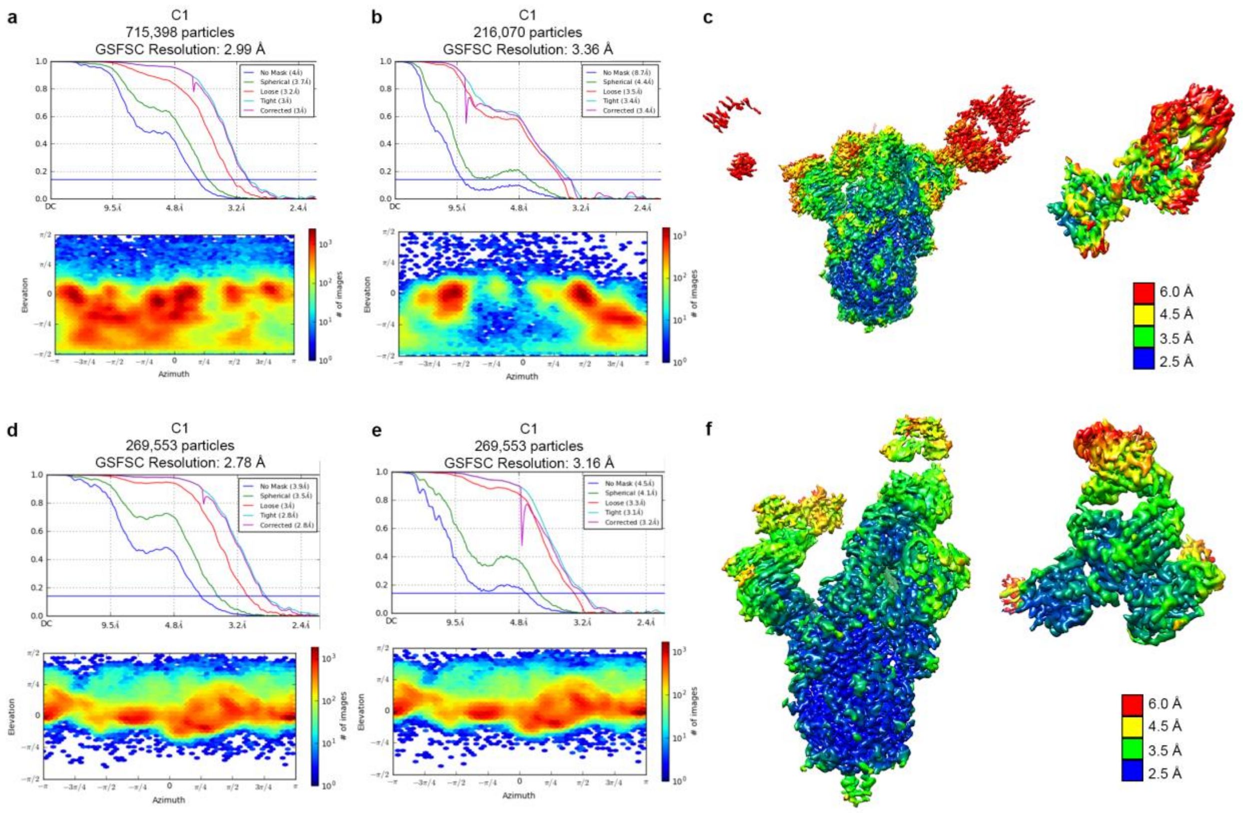


**Extended Figure 14. Cryo-EM data validation.**

**a-b,** FSC curves (top) and the viewing direction distribution plots (bottom) for global reconstruction of A7V3 bound to SARS-CoV-2 S [**a**] and focused reconstruction of A7V3 bound to S-NTD [**b**]. **c,** cryo-EM density map of A7V3 bound to SARS-CoV-2 S (left) and S-NTD (right), respectively. The local resolution is depicted by a spectrum of rainbow color as a scale bar. **d-e,** FSC curves (top) and the viewing direction distribution plots (bottom) for global reconstruction of N3-1 bound to SARS-CoV-2 S [**d**] and focused reconstruction of N3-1 bound to S-RBDs [**e**]. **f,** cryo-EM density map of N3-1 bound to SARS-CoV-2 S (left) and S-RBDs (right), respectively. The local resolution is depicted by a spectrum of rainbow color as a scale bar.

**Extended Data Table 3. N3-1 CryoEM Statistics**

| | | |
|---|---|---|
| **EM data collection** | | |
| Microscope | FEI Titan Krios | |
| Voltage (kV) | 300 | |
| Detector | Gatan K3 | |
| Magnification (nominal) | 22500 | |
| Pixel size (Å/pix) | 1.1 | |
| Flux ($e^-$/pix/sec) | 8 | |
| Frames per exposure | 80 | |
| Exposure ($e^-$/Å$^2$) | 80 | |
| Defocus range (μm) | 1.0-2.5 | |
| Micrographs collected | 3203 | |
| Sample | SARS-CoV-2 S + N3-1 Fab | |
| **3D reconstruction statistics** | | |
| | Overall | RBDs-N3-1 subcomplex |
| Particles | 269,553 | 269,553 |
| Symmetry | C1 | C1 |
| Map sharpening B-factor | -123.6 | -106.8 |
| Unmasked resolution at 0.143 FSC (Å) | 3.90 | 4.50 |
| Masked resolution at 0.143 FSC (Å) | 2.78 | 3.16 |
| **Model refinement and validation statistics** | | |
| Composition | | |
| Amino acids | | 579 |
| RMSD bonds (Å) | | 0.003 |
| RMSD angles (°) | | 0.59 |
| Average B-factors | | |
| Amino acids | | 102.3 |
| Ramachandran | | |
| Favored (%) | | 95.8 |
| Allowed (%) | | 4.2 |
| Outliers (%) | | 0 |
| Rotamer outliers (%) | | 0 |
| Clash score | | 4.0 |
| C-beta outliers (%) | | 0 |
| CaBLAM outliers (%) | | 1.6 |
| CC (mask) | | 0.83 |
| MolProbity score | | 1.48 |
| EMRinger score | | 4.95 |

**Extended Data Table 4. A7V3 CryoEM Statistics**

| **EM data collection** | | |
|---|---|---|
| Microscope | FEI Titan Krios | |
| Voltage (kV) | 300 | |
| Detector | Gatan K3 | |
| Magnification (nominal) | 22500 | |
| Pixel size (Å/pix) | 1.1 | |
| Flux ($e^{-}$/pix/sec) | 8 | |
| Frames per exposure | 80 | |
| Exposure ($e^{-}$/Å$^2$) | 80 | |
| Defocus range (μm) | 1.0-2.5 | |
| Micrographs collected | 3636 | |
| Sample | SARS-CoV-2 S + A7V3 Fab | |
| **3D reconstruction statistics** | | |
| | Overall | NTD-A7V3 subcomplex |
| Particles | 715,398 | 216,070 |
| Symmetry | C1 | C1 |
| Map sharpening B-factor | -151.7 | -94.1 |
| Unmasked resolution at 0.143 FSC (Å) | 4.00 | 8.70 |
| Masked resolution at 0.143 FSC (Å) | 2.99 | 3.36 |
| **Model refinement and validation statistics** | | |
| Composition | | |
| Amino acids | | 452 |
| RMSD bonds (Å) | | 0.003 |
| RMSD angles (º) | | 0.62 |
| Average B-factors | | |
| Amino acids | | 111.1 |
| Ramachandran | | |
| Favored (%) | | 94.0 |
| Allowed (%) | | 6.0 |
| Outliers (%) | | 0 |
| Rotamer outliers (%) | | 0 |
| Clash score | | 4.7 |
| C-beta outliers (%) | | 0 |
| CaBLAM outliers (%) | | 3.1 |
| CC (mask) | | 0.76 |
| MolProbity score | | 1.64 |
| EMRinger score | | 3.38 |

**Extended Data Table 5. Houston spike variant ELISAs**

| Variant | ACE2 | | CR3022 | | N3-1 | | A7V3 | |
|---|---|---|---|---|---|---|---|---|
| | EC50 nM | R squared | EC50 nM | R squared | EC50 nM | R squared | EC50 nM | R squared |
| **S-2P** | 8.5 | 0.99 | 3.6 | 0.99 | 0.8 | 0.96 | 1.8 | 0.97 |
| **S-2P D614G** | 7.4 | 0.95 | 3.9 | 0.98 | 1.4 | 0.99 | 4.1 | 0.98 |
| **HexaPro** | 8.9 | 0.95 | 7.3 | 0.97 | 1.6 | 0.99 | 8.7 | 0.99 |
| **HexaPro D614G** | 4.9 | 0.99 | 5.9 | 0.98 | 1.6 | 0.99 | 3.1 | 0.97 |
| **F338L** | 4.5 | 0.97 | 15.9 | 0.99 | 1.5 | 0.93 | 2.5 | 0.98 |
| **A352S** | 4.4 | 0.99 | 4.3 | 0.97 | 1.6 | 0.98 | 3.7 | 0.98 |
| **T385I** | 4.8 | 0.97 | 5.2 | 0.96 | 1.0 | 0.99 | 1.6 | 0.98 |
| **A419V** | 5.4 | 0.99 | 3.9 | 0.96 | 2.1 | 0.95 | 1.9 | 0.96 |
| **V445F** | 4.2 | 0.95 | 5.4 | 0.98 | 0.9 | 0.99 | 3.1 | 0.99 |
| **G446V** | 4.0 | 0.96 | 3.3 | 0.99 | 2.0 | 0.97 | 3.1 | 0.99 |
| **F456L** | 3.8 | 0.99 | 3.9 | 0.98 | 1.5 | 0.98 | 3.9 | 0.98 |
| **E484Q** | 4.0 | 0.96 | 3.9 | 0.98 | 2.1 | 0.99 | 2.6 | 0.98 |
| **A520S** | 3.8 | 0.96 | 2.4 | 0.98 | 1.3 | 0.99 | 4.5 | 0.97 |
| **K528R** | 5.4 | 0.99 | 3.6 | 0.98 | 1.0 | 0.96 | 5.7 | 0.97 |
| **S373P** | 3.8 | 0.95 | 93.1 | 0.97 | 3.7 | 0.99 | 1.7 | 0.98 |
| **R408T** | 8.2 | 0.96 | 10.1 | 0.99 | 1.5 | 0.99 | 10.4 | 0.94 |

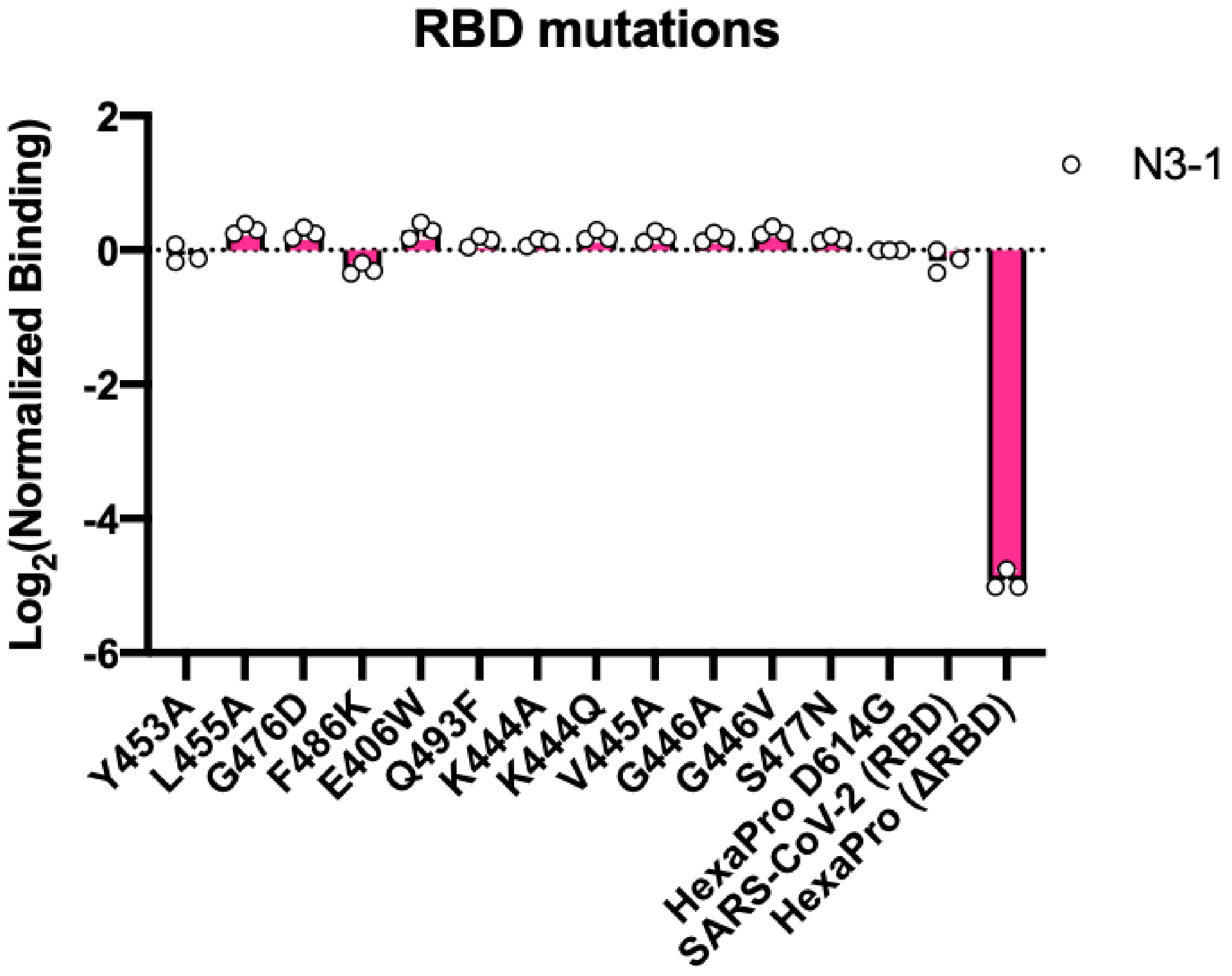


**Extended Figure 15. N3-1 binding to Regeneron escape mutants and S477N, as measured using the mammalian surface display assay.** HEK293T cells transiently expressing full length spike protein were stained with anti-spike antibodies and analyzed by flow cytometry. The median fluorescence intensity of the stained cells was normalized to the HexaPro-D614G spike. Spike variants with single RBD mutations shown to reduce REGN10987 and/or REGN10933 binding were tested with N3-1. The SARS-CoV2 RBD subunit and SARS-CoV-2 spike with a deleted RBD (ΔRBD) were included as controls.
